# Sensory neuron LKB1 mediates ovarian and reproductive function

**DOI:** 10.1101/2023.03.28.534533

**Authors:** Melissa E Lenert, Michael D Burton

**Author notes:** Corresponding Author: Michael D Burton, PhD, The University of Texas at Dallas, 800 W Campbell Rd, Richardson TX 75080.

## Abstract

Treatments for reproductive disorders in women primarily consist of hormone replacement therapy, which can have negative health impacts. Bidirectional communication between sensory neurons and innervated organs is an emerging area of interest in tissue physiology with potential relevance for reproductive disorders. Indeed, the metabolic activity of sensory neurons can have profound effects on reproductive phenotypes. To investigate this phenomenon, we utilized a murine model with conditional deletion in sensory neurons of liver kinase B1 (LKB1), a serine/threonine kinase that regulates cellular metabolism. Female mice with this LKB1 deletion (Na_v_1.8cre;LKB1^fl/fl^) had significantly more pups per litter compared to wild-type females. Interestingly, the LKB1 genotype of male breeders had no effect on fertility outcomes, thus indicating a female-specific role of sensory neuron metabolism in fertility. LKB1 deletion in sensory neurons resulted in reduced ovarian innervation from dorsal root ganglia neurons and increased follicular turnover compared to littermate controls. In summary, LKB1 expression in peripheral sensory neurons plays an important role in modulating fertility of female mice via ovarian sensory innervation.

## Introduction

Infertility and disorders of the reproductive system affect around 12% of women of reproductive age (Agarwal *et al*, 2015; Carson & Kallen, 2021). While both men and women can experience infertility, ovarian dysfunction contributes to 25% of infertility in couples (Carson & Kallen, 2021). The primary factors that impair female fertility include metabolic stress, hormonal imbalance, and increasing age (Christensen *et al*, 2012; Colledge, 2013; Mikhael *et al*, 2019). The mammalian ovary is the primary site of female fertility, being the major producer of female sex hormones and to locus for preparation of oocytes for fertilization via follicle growth and ultimately ovulation (Fauser & van Heusden, 1997; Hsueh *et al*, 2015). Ovulation is a highly regulated process that is dependent on coordinated shifts in sex hormones levels, oocyte maturation, and paracrine communication within the ovary (Gershon & Dekel, 2020; Robker *et al*, 2018). Whether the patient’s goal is to promote ovulation or prevent it, the first line treatment entails administration of exogenous hormones, which often have poorly tolerated off-target adverse effects (Brabaharan *et al*, 2022; Engler-Chiurazzi *et al*, 2022; Mørch *et al*, 2017; Robakis *et al*, 2019). Thus, identifying non-hormonal processes involved in the regulation of fertility is a matter of high importance.

There is a well-established relationship between cellular metabolism and reproductive fitness: metabolic stress has profoundly detrimental effects on fertility. States of perturbed metabolism due to conditions such as obesity or prolonged malnutrition result in impaired ovulation and steroid hormone production (Deng *et al*, 2017; Lainez & Coss, 2019; Magyar *et al*, 2022; Sirotkin *et al*, 2018; Wassif *et al*, 2011). This synergy of metabolism and reproductive function may arise from effects of the liver kinase B1 (LKB1) gene, a ubiquitously expressed serine/threonine kinase with multiple downstream targets involved in metabolism, cellular growth, and mitochondrial homeostasis (Molina *et al*, 2021; Nakano & Takashima, 2012; Shackelford & Shaw, 2009). There is extensive research of LKB1-dependent pathways in cancer and metabolic disorders, but relatively little knowledge about its role in modulating peripheral nervous system function (Alessi *et al*, 2006; Herzig & Shaw, 2018; Saikia & Joseph, 2021). We have recently used a novel mouse model with sensory neuron specific deletion of LKB1 to show that metabolism in sensory neurons is an important regulator of physiological responses to fasting in female mice(Garner & Burton, 2022). Sensory neurons in female mice with conditional knockout of LKB1 had significantly diminished metabolic activity after fasting compared to wild-type (WT) controls (Garner & Burton, 2022). Interestingly, metabolic activity in male sensory neurons was not LKB1-dependent, suggesting a female-specific role of LKB1 signaling in murine sensory neurons.

Several studies in recent years have demonstrated that communication between peripheral sensory neurons and the organs they innervate influences the functional activity of recipient organs. This phenomenon occurs in multiple tissue types, including the lymph nodes, adipose tissue, the gut, and dura mater of the brain (Chiu *et al*, 2013; Huang *et al*, 2021; Wang *et al*, 2022). Sensory neurons regulate aspects of local microenvironment at the site of innervation by detecting and responding to noxious and innocuous stimuli such as inflammation, metabolic state, tissue remodeling, and hormones (Lenert *et al*, 2021a; Pinho-Ribeiro *et al*, 2017). Key studies in the present context have demonstrated an important role of peripheral sensory neurons in ovarian hormone output and ovulation (Flores *et al*, 2011; Kagitani *et al*, 2008; Morales-Ledesma *et al*, 2010; Ramírez Hernández *et al*, 2020; Rosas *et al*, 2018). The ovaries are innervated by sympathetic and sensory ganglia (Burden *et al*, 1983; Calka *et al*, 1988; Lawrence Jr & Burden, 1980; Pastelín *et al*, 2017). Signaling via sympathetic innervation (primarily tyrosine hydroxylase expressing) influences hormone release, folliculogenesis, and ovulation; however, the role of sensory neurons in these processes is unclear (Cruz *et al*, 2017; Morales-Ledesma *et al*, 2015; Zoubina & Smith, 2001). Several research groups have shown that the dorsal root ganglia (DRG), which houses the cell bodies of peripheral sensory neurons, innervate the ovary in healthy female rats (Burden *et al*., 1983; Inyama *et al*, 1986). Dysregulation in sympathetic neuron signaling and innervation in the ovary has been reported in human fertility disorders and in corresponding animal models such as polycystic ovarian syndrome and endometriosis; a few studies suggest that an imbalance between sensory and sympathetic signaling/innervation to the reproductive tract is part of their pathogenesis (Arnold *et al*, 2012; del Campo *et al*, 2019; Lara *et al*, 2002; Morales-Ledesma *et al*., 2010). Obtaining a better understanding of the role of sensory neuron metabolism and innervation in the ovary should open new therapeutic channels for the treatment of fertility disorders.

The aim of the current study is to characterize the effects of LKB1 deletion in murine peripheral sensory neurons on fertility outcomes, ovarian growth and turnover, and innervation. We had observed that our mouse line with LKB1-null sensory neurons were prolific breeders. Female breeders with LKB1-null sensory neurons had much larger litter sizes than their WT counterparts, independent of sensory neuron LKB1-deletion in male breeders. To determine the effects of sensory neuron LKB1-deletion on reproductive phenotypes, we assessed breeding metrics, estrous cycling, and serum estradiol levels. We then hypothesized that females with LKB1-null sensory neurons would have reduced sensory innervation in the ovary and an increased number of viable follicles compared to WT littermate controls. To understand the role of LKB1 in ovarian innervation, we performed retrograde tract tracing of ovarian innervation by DRG neurons. To further explore the mechanism of LKB1-deletion effects on ovulation and fertility, we measured follicle growth and development in the ovaries. In summary, LKB1 deletion from sensory neurons enhances fertility in female mice and presents a novel target for future development of fertility therapeutics.

## Results

### Female mice with LKB1-null sensory neurons have enhanced fertility

We had early anecdotal observations that female breeders with LKB1-null sensory neurons (Na_v_1.8^LKB1^) had larger litters than WT breeders. Our breeding schema was such that male and female mice have opposite genotypes, i.e., LKB1^fl/fl^ (LKB1^fl/fl^, normal expression of LKB1) or Na_v_1.8^LKB1^. To assess whether the increased litter sizes were due to the reproductive phenotype of the male or female breeders, we compared the breeding histories of 1) Na_v_1.8^LKB1^ females paired with LKB1^fl/fl^ males, 2) Na_v_1.8^LKB1^ males paired with LKB1^fl/fl^ females, and 3) WT males paired with WT females (Figure 1A, Table 1). When the breeding pairs included a Na_v_1.8^LKB1^ females, the number of pups per litter (Na_v_1.8^LKB1^: mean ± SEM = 12.1 ± 0.8) was significantly greater than when the breeding pairs had a LKB1^fl/fl^ female (LKB1^fl/fl^: mean ± SEM = 7.4 ± 0.45) (Figure 1B-C, Table 4). Further, LKB1^fl/fl^ females paired with Na_v_1.8^LKB1^ males produced similar litter sizes compared to WT x WT pairings (WT: mean ± SEM = 5.7 ± 0.47), indicating that deletion of LKB1 from peripheral sensory neurons in males had no direct effect on litter size. We also compared litter sizes from WT mice of other inbred strains and the outbred ICR strain (Supplemental Figure 1C, Supplemental Table 1). There were no differences in litter sizes between any of the WT mixed strain breeders, whereas outbred ICR breeders had significantly larger litter sizes than WT inbred mice. Litter sizes for Na_v_1.8^LKB1^ and ICR female breeders were similar. Because we observed larger litter sizes only with Na_v_1.8^LKB1^ female/LKB1^fl/fl^ male breeding pairs, it appears that LKB1 expression in sensory neurons plays a female-specific role in modulating fertility.

**Figure 1.**
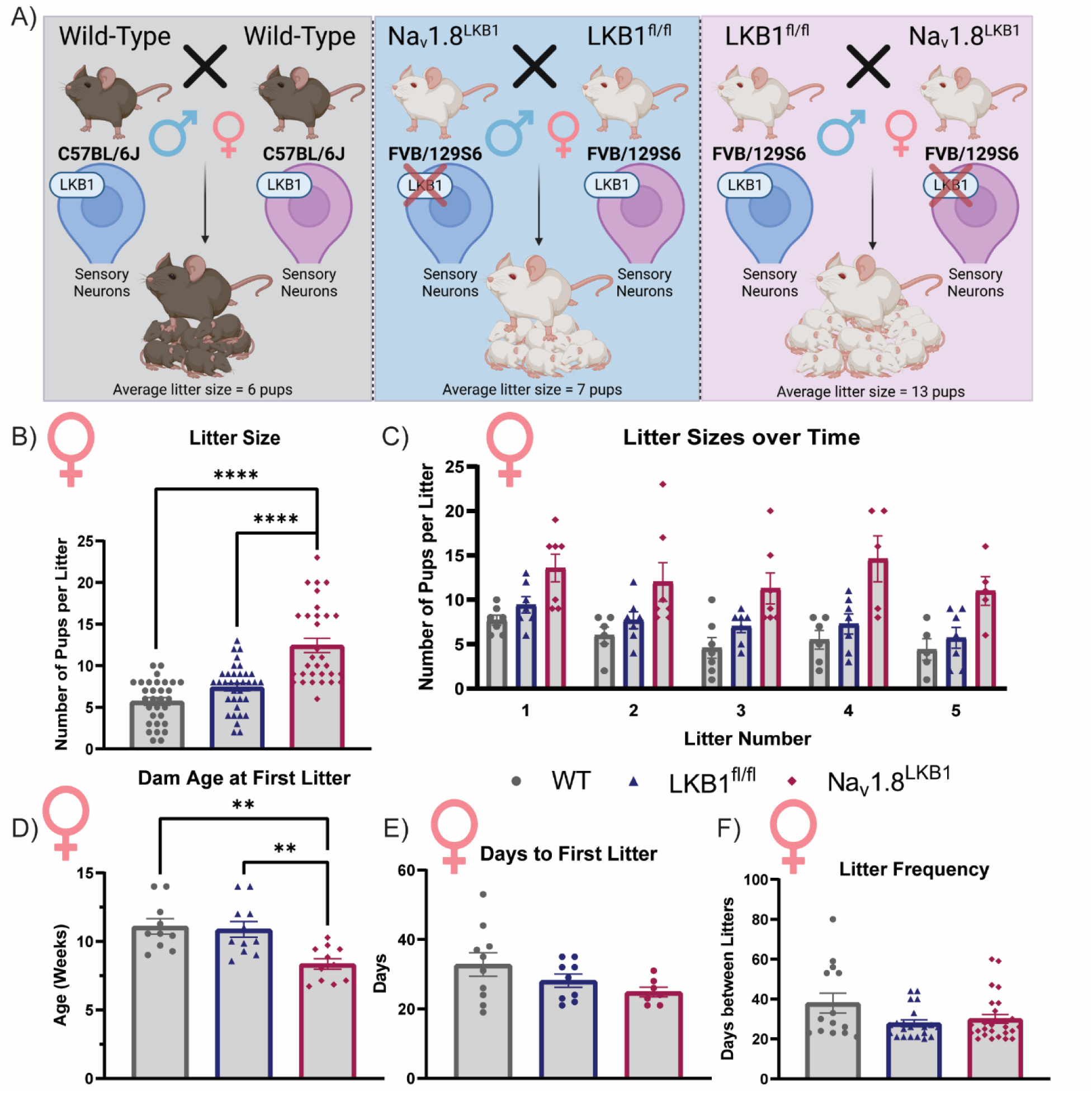
Female mice with LKB1-null sensory neurons have enhanced fertility. (A) Graphic depiction of our breeding schema. (B) Number of pups per litter (WT: mean ± SEM = 5.7 ± 0.47 viable pups, n=31 litters from 10 females with 2-5 litters each; LKB1^fl/fl^: mean ± SEM = 7.4 ± 0.45 viable pups, n=35 litters from 11 females with 2-5 litters each; Na_v_1.8^LKB1^: mean ± SEM = 12.1 ± 0.8 viable pups, n=31 litters from 11 females with 2-5 litters each). (C) Number of pups per litter per female breeder across the first five litters. (D) Dam age in weeks at the birth of their first litter (WT: mean ± SEM = 11.1 ± 5.6 weeks, n=10 females; LKB1^fl/fl^: mean ± SEM = 10.9 ± 0.57 weeks, n=11 females; Na_v_1.8^LKB1^: mean ± SEM = 8.4± 0.38 weeks, n=11 females). (E) Number of days between breeding initiation and the birth of their first litter (WT: mean ± SEM = 32.8 ± 3.4 days, n = 10 females; LKB1^fl/fl^: mean ± SEM = 28.1 ± 1.9 days, n=9 females; Na_v_1.8^LKB1^: mean ± SEM = 24.9 ± 1.4 days, n=7 females). (F) Number of days between consecutive litters per female breeder (WT: mean ± SEM = 38.0 ± 4.96 days, n=14 from 10 females with 2-5 litters each; LKB1^fl/fl^: mean ± SEM = 27.9 ± 1.7 days, n=21 from 11 females with 2-5 litters each; Na_v_1.8^LKB1^: mean ± SEM = 30.0 ± 2.2, n=26 from 11 females with 2-5 litters each). *p<0.05, **p<0.01, ***p<0.0001, ****p<0.0001 by ordinary one-way ANOVA (B, D, E, and F) or repeated measures two-way ANOVA (C).

**Table 1.** Breeding Schema and Metrics. All values are represented as mean ± SEM.

| Strain Composition | Dam Genotype | Sire Genotype | Number of Dams | Dam Age (Weeks) at First Litter | Days to First Litter | Litter Size | Days between Consecutive Litters |
| --- | --- | --- | --- | --- | --- | --- | --- |
| C57BL/6J | Wild-Type | Wild-Type | 10 | 11.10 ± 0.5638 | 32.80 ± 3.399 | 5.710 ± 0.4711 | 38.00 ± 4.957 |
| FVB, 129S6, C57BL/6J | LKB1<sup>fl/fl</sup> | Na<sub>v</sub>1.8<sup>LKB1</sup> | 11 | 10.88 ± 0.5720 | 28.11 ± 1.889 | 7.429 ± 2.747 | 27.86 ± 1.695 |
| FVB, 129S6, C57BL/6J | Na<sub>v</sub>1.8<sup>LKB1</sup> | LKB1<sup>fl/fl</sup> | 11 | 8.351 ± 0.3830 | 24.86 ± 1.405 | 12.45 ± 0.8486 | 30.04 ± 2.240 |
| FVB, 129S6, C57BL/6J | Wild-Type | Wild-Type | 6 | 14.05 ± 2.148 | 28.83 ± 2.455 | 7.222 ± 0.4754 | 33.14 ± 3.477 |
| ICR (outbred) | Wild-Type | Wild-Type | 5 | 10.34 ± 0.6037 | 24.0 ± 0.9487 | 13.28 ± 1.315 | 59.62 ± 6.075 |

To further understand the reproductive phenotype of Na_v_1.8^LKB1^ females, we determined the age at which female breeders had their first litter. All breeders were paired between 5-6 weeks of age. Na_v_1.8^LKB1^ females had their first litter at a significantly younger age than either LKB1^fl/fl^ mice or WT mice of other strains (Figure 1D, Table 2; Supplemental Figure 1D, Supplemental Table 1). Further, there were no significant differences in the number of days between breeding initiation and the first litter birth (Figure 1E, Table 2; Supplemental Figure 1E, Supplemental Table 1). Thus, we attribute the younger age of first pregnancy to precocious puberty observed in Na_v_1.8^LKB1^ females. We also assessed litter frequency (as mean days between the birth of consecutive litters from each female) and found no significant differences in the frequency of litters between Na_v_1.8^LKB1^, LKB1^fl/fl^, and WT females (Figure 1F, Table 2, Supplemental Figure 1F, Supplemental Table 1). Taken together, these results show that LKB1 in sensory neurons modulates the onset of reproductive maturity and increases litter sizes, but likely does not play a significant role in the frequency of pregnancy in female mice. The presence of LKB1 in sensory neurons in male mice does not seem to affect their reproductive phenotype. Thus, these data support the novel finding that LKB1 expression in sensory neurons modulates fertility and reproductive physiology in a female-specific manner.

**Table 2.** Statistics corresponding to Figure 1. Litter sizes, age at first litter, days to first litter, and litter frequency were analyzed using ordinary one-way ANOVA with post hoc Tukey’s multiple comparisons. Litter sizes over time was analyzed using repeated measures two-way ANOVA with post hoc Tukey’s multiple comparisons between genotypes at each litter. Significance was set at p<0.05 for all datasets. All significant results are in bold.

| Figure | Dataset | Main Effect |  | Multiple Comparisons |  |
| --- | --- | --- | --- | --- | --- |
|  |  | F (DFn, DFd) | p-value | Groups | p-value |
| 1B | Litter Sizes | F (2, 94) = 31.77 | **p<0.0001** | WT vs LKB1<sup>fl/fl</sup><br>**WT vs Na<sub>v</sub>1.8<sup>LKB1</sup>**<br>LKB1<sup>fl/fl</sup> vs Na<sub>v</sub>1.8<sup>LKB1</sup> | p=0.1159<br>**p<0.0001**<br>**p<0.0001** |
| 1C | Litter Sizes Over Time | Litter: F (2.707, 43.31) = 3.683<br>Genotype: F (2, 18) = 15.63<br>Litter x Genotype: F (8, 64) = 0.3204 | **p=0.0223**<br>**p=0.0001**<br>p=0.9554 | Litter 1: WT vs LKB1<sup>fl/fl</sup><br>**Litter 1: WT vs Na<sub>v</sub>1.8<sup>LKB1</sup>**<br>Litter 1: LKB1<sup>fl/fl</sup> vs Na<sub>v</sub>1.8<sup>LKB1</sup><br>Litter 2: WT vs LKB1<sup>fl/fl</sup><br>Litter 2: WT vs Na<sub>v</sub>1.8<sup>LKB1</sup><br>Litter 2: LKB1<sup>fl/fl</sup> vs Na<sub>v</sub>1.8<sup>LKB1</sup><br>Litter 3: WT vs LKB1<sup>fl/fl</sup><br>**Litter 3: WT vs Na<sub>v</sub>1.8<sup>LKB1</sup>**<br>Litter 3: LKB1<sup>fl/fl</sup> vs Na<sub>v</sub>1.8<sup>LKB1</sup><br>Litter 4: WT vs LKB1<sup>fl/fl</sup><br>**Litter 4: WT vs Na<sub>v</sub>1.8<sup>LKB1</sup>**<br>Litter 4: LKB1<sup>fl/fl</sup> vs Na<sub>v</sub>1.8<sup>LKB1</sup><br>Litter 5: WT vs LKB1<sup>fl/fl</sup><br>**Litter 5: WT vs Na<sub>v</sub>1.8<sup>LKB1</sup>**<br>Litter 5: LKB1<sup>fl/fl</sup> vs Na<sub>v</sub>1.8<sup>LKB1</sup> | p=0.2963<br>**p=0.0203**<br>p=0.1047<br>p=0.4278<br>p=0.0807<br>p=0.2280<br>p=0.2248<br>**p=0.0224**<br>p=0.1166<br>p=0.5085<br>**p=0.0469**<br>p=0.0982<br>p=0.7226<br>**p=0.0298**<br>p=0.0683 |
| 1D | Age at First Litter | F (2, 29) = 9.040 | **p=0.0009** | WT vs LKB1<sup>fl/fl</sup><br>**WT vs Na<sub>v</sub>1.8<sup>LKB1</sup>**<br>LKB1<sup>fl/fl</sup> vs Na<sub>v</sub>1.8<sup>LKB1</sup> | p=0.9537<br>**p=0.0021**<br>**p=0.0036** |
| 1E | Days to First Litter | F (2, 23) = 2.265 | p=0.1266 | n/a |  |
| 1F | Litter Frequency | F (2, 58) = 2.974 | p=0.0590 | n/a |  |

Fertility and successful pregnancies decline with increasing age of women and female animals. Therefore, to determine if the enhanced fertility observed in Na_v_1.8^LKB1^ females is age-dependent, we assessed litter sizes in a small cohort of Na_v_1.8^LKB1^ and LKB1^fl/fl^ breeders at 20 weeks of age at the time of breeding initiation (Supplementary Figure 1G, Supplemental Table 1). All other parameters (breeding schema, housing conditions, handling) were matched to those used for young breeders. As expected, there was an age-dependent decrease in the number of pups per litter for both genotypes; however, there were no differences in the mean number of pups per litter between LKB1^fl/fl^ (mean ± SEM = 5.7 ± 1.0) and Na_v_1.8^LKB1^ (mean ± SEM = 7.4 ± 0.85) female breeders. These data suggest that the presence of LKB1 in sensory neurons in female mice stimulates follicle use and rates of ovulation to a decreasing extent with increasing maternal age.

### Females with LKB1-null sensory neurons have altered estrous cycling and lower serum estradiol

To investigate female-specific sources of variation in fertility, we assessed the estrous cycling patterns and serum estradiol levels in WT, LKB1^fl/fl^, and Na_v_1.8^LKB1^ females. We obtained vaginal lavage samples twice per day for ten days, thus capturing two typical murine estrous cycles of 4-5 days. To test for an association between altered breeding rate and shifts in estrous cycling, we analyzed the amount of time the animals were spending in each phase. Compared to LKB1^fl/fl^ females, Na_v_1.8^LKB1^ females spent less time in proestrus, whereas WT and LKB1^fl/fl^ mice had similar proestrus duration (Figure 2A, Table 3). LKB1^fl/fl^ and Na_v_1.8^LKB1^ females both spent more time in estrus than WT females, but without any group differences in metestrus or diestrus duration (Figure 2B-D, Table 3). Additionally, we did not observe any differences in cycle length according to strain or genotype (Figure 2E, Table 3). Measurements of serum estradiol levels across the estrus cycle indicated that WT and Na_v_1.8^LKB1^ females had lower estradiol than LKB1^fl/fl^ littermates during diestrus (Figure 2F, Table 3).

**Figure 2.**
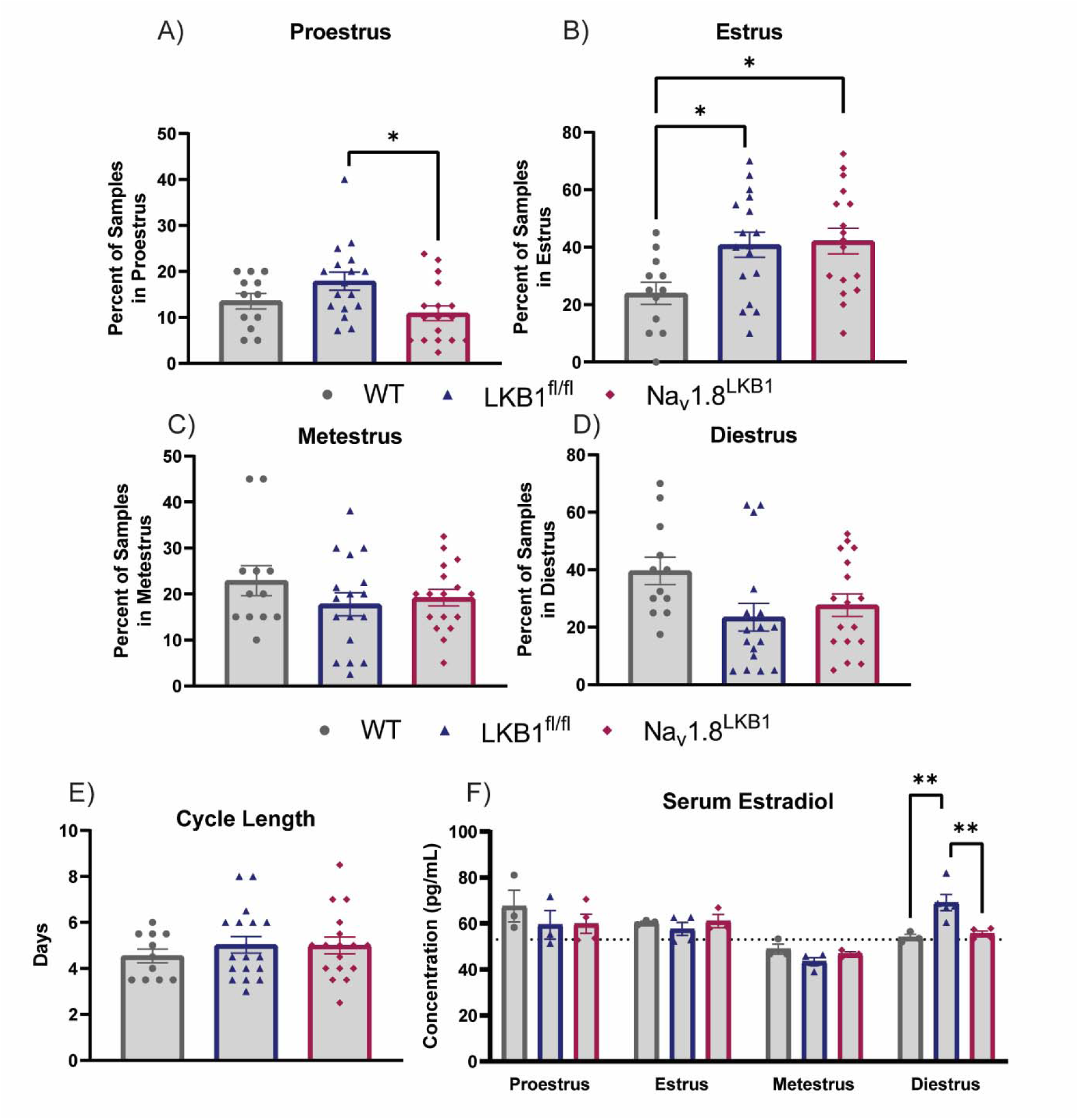
Females with LKB1-null sensory neurons have altered estrous cycling and lower serum estradiol levels. Estrous cycling was tracked for ten days with twice daily vaginal lavage sample collection. The percentage of samples for each mouse in (A) proestrus, (B) estrus, (C), metestrus, and (D) diestrus across the entire collection period. (A) Proestrus: (WT: mean ± SEM = 13.5 ± 1.7%; LKB1^fl/fl^: mean ± SEM = 17.9 ± 1.97%; Na_v_1.8^LKB1^: mean ± SEM = 10.9 ± 1.6%. (B) Estrus: (WT: mean ± SEM = 23.96 ± 3.8%; LKB1^fl/fl^: mean ± SEM = 40.8 ± 4.3%; Na_v_1.8^LKB1^: mean ± SEM = 42.2 ± 4.5%. (C) Metestrus: WT: mean ± SEM = 22.9 ± 3.3%; LKB1^fl/fl^: mean ± SEM = 17.8 ± 2.5%; Na_v_1.8^LKB1^: mean ± SEM = 19.2 ± 1.8%. (D) Diestrus: WT: mean ± SEM = 39.6 ± 4.8%; LKB1^fl/fl^: mean ± SEM = 23.5± 4.8%; Na_v_1.8^LKB1^: mean ± SEM = 27.7 ± 3.9%. (E) Total cycle length (days between proestrus smears): (WT: mean ± SEM = 4.5 ± 0.27 days; LKB1^fl/fl^: mean ± SEM = 4.9 ± 0.33 days; Na_v_1.8^LKB1^: mean ± SEM = 5.0 ± 0.36 days. For (A-E), WT: n=12 females; LKB1^fl/fl^: n=17 females; Na_v_1.8^LKB1^: n=17 females. (F) Serum estradiol (pg/mL) levels across the estrous cycle. (WT: n=3 biological replicates per phase; LKB1^fl/fl^: n=3-5 biological replicates per phase; Na_v_1.8^LKB1^: n=3-5 biological replates per phase; all biological replicates were assayed in duplicate). The dotted line indicates the mean serum estradiol (53 pg/mL) for age-matched male mice. *p<0.05, **p<0.01, ***p<0.0001, ****p<0.0001 by ordinary one-way ANOVA (A-E) or ordinary two-way ANOVA (F).

**Table 3.** Statistics corresponding to Figure 2. Percentage of samples in proestrus, estrus, metestrus, and diestrus as well as cycle length were analyzed using ordinary one-way ANOVA with *post hoc* Tukey’s multiple comparisons. Serum estradiol was analyzed using ordinary two-way ANOVA with *post hoc* Tukey’s correction for multiple comparisons between genotypes at each phase. Significance was set at *p*<0.05 for all datasets. All significant results are in bold.

| Figure | Dataset | Main Effect |  | Multiple Comparisons |  |
| --- | --- | --- | --- | --- | --- |
|  |  | F (DFn, DFd) | p-value | Groups | p-value |
| 2A | Proestrus | F (2, 43) = 4.248 | **p=0.0207** | WT vs LKB1<sup>fl/fl</sup><br>WT vs Na<sub>v</sub>1.8<sup>LKB1</sup><br>LKB1<sup>fl/fl</sup> vs Na<sub>v</sub>1.8<sup>LKB1</sup> | p=0.2386<br>p=0.5894<br>**p=0.0162** |
| 2B | Estrus | F (2, 43) = 4.731 | **p=0.0139** | WT vs LKB1<sup>fl/fl</sup><br>WT vs Na<sub>v</sub>1.8<sup>LKB1</sup><br>LKB1<sup>fl/fl</sup> vs Na<sub>v</sub>1.8<sup>LKB1</sup> | **p=0.0312**<br>**p=0.0187**<br>p=0.9718 |
| 2C | Metestrus | F (2, 43) = 1.035 | p=0.3638 | n/a |  |
| 2D | Diestrus | F (2, 43) = 2.999 | p=0.0603 | n/a |  |
| 2E | Cycle Length | F (2, 43) = 0.4728 | p=0.6265 | n/a |  |
| 2F | Serum Estradiol | Phase: F (3, 30) = 12.49<br>Genotype: F (2, 30) = 0.3404<br>Phase x Genotype: F (6, 30) = 3.021 | **p<0.0001**<br>p=0.7142<br>**p=0.0197** | Proestrus: WT vs LKB1<sup>fl/fl</sup><br>Proestrus: WT vs Na<sub>v</sub>1.8<sup>LKB1</sup><br>Proestrus: LKB1<sup>fl/fl</sup> vs Na<sub>v</sub>1.8<sup>LKB1</sup><br>Estrus: WT vs LKB1<sup>fl/fl</sup><br>Estrus: WT vs Na<sub>v</sub>1.8<sup>LKB1</sup><br>Estrus: LKB1<sup>fl/fl</sup> vs Na<sub>v</sub>1.8<sup>LKB1</sup><br>Metestrus: WT vs LKB1<sup>fl/fl</sup><br>Metestrus: WT vs Na<sub>v</sub>1.8<sup>LKB1</sup><br>Metestrus: LKB1<sup>fl/fl</sup> vs Na<sub>v</sub>1.8<sup>LKB1</sup><br>**Diestrus: WT vs LKB1<sup>fl/fl</sup>**<br>Diestrus: WT vs Na<sub>v</sub>1.8<sup>LKB1</sup><br>**Diestrus: LKB1<sup>fl/fl</sup> vs Na<sub>v</sub>1.8<sup>LKB1</sup>** | p=0.2830<br>p=0.2752<br>p=0.9956<br>p=0.8260<br>p=0.9933<br>p=0.7601<br>p=0.5127<br>p=0.9137<br>p=0.7761<br>**p=0.0082**<br>p=0.9455<br>**p=0.0099** |

### LKB1 in peripheral sensory neurons promotes ovarian innervation

To investigate whether Na_v_1.8 innervation originated from DRGs, we performed retrograde tracing from the ovary in WT, LKB1^fl/fl^, and Na_v_1.8^LKB1^ mice by injecting with Fluorogold and collecting DRGs ten days later at both thoracic (T7-T13) and lumbar (L1-L5) levels of the spinal cord (Figure 3A). Assessment of the total number of fluorogold-labeled neurons per DRG in WT C57BL/6J mice showing fluorogold labelling in thoracic (T10-T13, especially T13) and lumbar (L1) DRGs (Figure 3B), but no labelling outside those spinal levels. These findings are congruent with earlier results in rats showing ovarian innervation arises from DRG extending from levels T10-L2 (Burden *et al*., 1983). WT mice had greater innervation than LKB1^fl/fl^ mice from T12-L1, but comparable fluorogold labelling staining at T10-T11. In particular, female Na_v_1.8^LKB1^ mice had relatively low fluorogold staining in DRG at levels T12-T13 and L1, with no differences in fluorogold staining in T10-T11 compared to WT (Figure 3B, Table 4).

**Figure 3.**
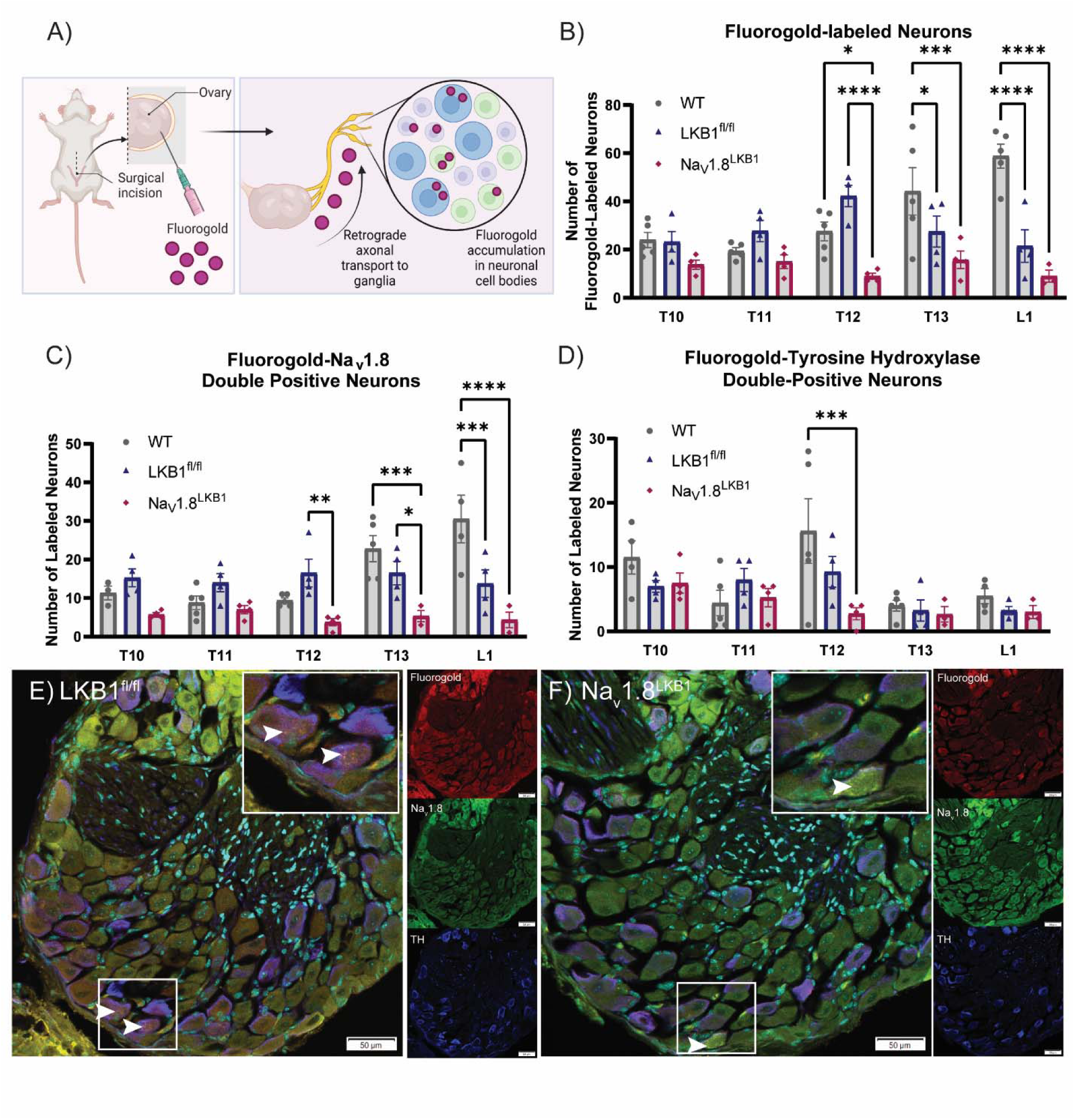
LKB1 in peripheral sensory neurons promotes ovarian innervation. (A) Graphic schema of the experimental methods used for retrograde tracing of ovarian innervation. (B) The number of fluorogold-positive neurons per DRG. (WT: n=5 mice; LKB1^fl/fl^: n=4 mice, Na_v_1.8^LKB1^: n=4 mice.) For each group, we obtained n=5 technical replicates per DRG. (C) The number of Na_v_1.8-positive neurons with positive fluorogold signal per DRG. (WT: n=5 mice; LKB1^fl/fl^: n=4 mice, Na_v_1.8^LKB1^: n=4 mice.) For each group, we obtained n=5 technical replicates per DRG. (D) The number of TH-positive neurons with positive fluorogold signal per DRG. (WT: n=5 mice; LKB1^fl/fl^: n=4 mice, Na_v_1.8^LKB1^: n=4 mice.) For each group, we used n=5 technical replicates per DRG. (E-F) Representative confocal microscopy (30X) of fluorogold (red), Na_v_1.8 (green), and TH (purple) immunohistochemistry in the T12 DRG from LKB1^fl/fl^ (E) and Na_v_1.8^LKB1^ (F) females. Scale bar = 50 µm. Arrows indicate positive fluorogold signal. *p<0.05, **p<0.01, ***p<0.0001, ****p<0.0001 by repeated measures two-way ANOVA (D-F).

**Table 4.** Statistics corresponding to Figure 3. Fluorogold expression, Na_v_1.8, and TH colocalization were analyzed using ordinary two-way ANOVA with post hoc Tukey’s multiple comparisons to perform genotype comparisons at each DRG. Significance was set at p<0.05 for all datasets. All significant results are in bold.

| Figure | Dataset | Main Effect |  | Multiple Comparisons |  |
| --- | --- | --- | --- | --- | --- |
|  |  | F (DFn, DFd) | p-value | Groups | p-value |
| 3E | Fluorogold (+) Neurons | DRG: F (4, 49) = 2.459<br>Genotype: F (2, 49) = 26.42<br>DRG x Genotype: F (8, 49) = 6.066 | $p=0.0576$<br>**$p<0.0001$**<br>**$p<0.0001$** | T10: WT vs LKB1<sup>fl/fl</sup><br>T10: WT vs Na<sub>v</sub>1.8<sup>LKB1</sup><br>T10: LKB1<sup>fl/fl</sup> vs Na<sub>v</sub>1.8<sup>LKB1</sup><br>T11: WT vs LKB1<sup>fl/fl</sup><br>T11: WT vs Na<sub>v</sub>1.8<sup>LKB1</sup><br>T11: LKB1<sup>fl/fl</sup> vs Na<sub>v</sub>1.8<sup>LKB1</sup><br>T12: WT vs LKB1<sup>fl/fl</sup><br>**T12: WT vs Na<sub>v</sub>1.8<sup>LKB1</sup>**<br>**T12: LKB1<sup>fl/fl</sup> vs Na<sub>v</sub>1.8<sup>LKB1</sup>**<br>**T13: WT vs LKB1<sup>fl/fl</sup>**<br>**T13: WT vs Na<sub>v</sub>1.8<sup>LKB1</sup>**<br>T13: LKB1<sup>fl/fl</sup> vs Na<sub>v</sub>1.8<sup>LKB1</sup><br>**L1: WT vs LKB1<sup>fl/fl</sup>**<br>**L1: WT vs Na<sub>v</sub>1.8<sup>LKB1</sup>**<br>L1: LKB1<sup>fl/fl</sup> vs Na<sub>v</sub>1.8<sup>LKB1</sup> | $p=0.9934$<br>$p=0.2989$<br>$p=0.3907$<br>$p=0.4452$<br>$p=0.7962$<br>$p=0.1895$<br>$p=0.0910$<br>**$p=0.0237$**<br>**$p<0.0001$**<br>**$p=0.0466$**<br>**$p=0.0004$**<br>$p=0.2415$<br>**$p<0.0001$**<br>**$p<0.0001$**<br>$p=0.2516$ |
| 3F | Fluorogold & Na<sub>v</sub>1.8 (+) Neurons | DRG: F (4, 45) = 3.329<br>Genotype: F (2, 45) = 23.05<br>DRG x Genotype: F (8, 45) = 4.545 | **$p=0.0180$**<br>**$p<0.0001$**<br>**$p=0.0004$** | T10: WT vs LKB1<sup>fl/fl</sup><br>T10: WT vs Na<sub>v</sub>1.8<sup>LKB1</sup><br>T10: LKB1<sup>fl/fl</sup> vs Na<sub>v</sub>1.8<sup>LKB1</sup><br>T11: WT vs LKB1<sup>fl/fl</sup><br>T11: WT vs Na<sub>v</sub>1.8<sup>LKB1</sup><br>T11: LKB1<sup>fl/fl</sup> vs Na<sub>v</sub>1.8<sup>LKB1</sup><br>T12: WT vs LKB1<sup>fl/fl</sup><br>T12: WT vs Na<sub>v</sub>1.8<sup>LKB1</sup><br>**T12: LKB1<sup>fl/fl</sup> vs Na<sub>v</sub>1.8<sup>LKB1</sup>**<br>T13: WT vs LKB1<sup>fl/fl</sup><br>**T13: WT vs Na<sub>v</sub>1.8<sup>LKB1</sup>**<br>**T13: LKB1<sup>fl/fl</sup> vs Na<sub>v</sub>1.8<sup>LKB1</sup>**<br>**L1: WT vs LKB1<sup>fl/fl</sup>**<br>**L1: WT vs Na<sub>v</sub>1.8<sup>LKB1</sup>**<br>L1: LKB1<sup>fl/fl</sup> vs Na<sub>v</sub>1.8<sup>LKB1</sup> | $p=0.6291$<br>$p=0.3944$<br>$p=0.0508$<br>$p=0.3542$<br>$p=0.8798$<br>$p=0.1876$<br>$p=0.1493$<br>$p=0.2936$<br>**$p=0.0062$**<br>$p=0.2204$<br>**$p=0.0003$**<br>**$p=0.0307$**<br>**$p=0.0003$**<br>**$p<0.0001$**<br>$p=0.0790$ |
| 3G | Fluorogold & TH (+) Neurons | DRG: F (4, 46) = 4.211<br>Genotype: F (2, 46) = 3.883<br>DRG x Genotype: F (8, 46) = 1.820 | **$p=0.0055$**<br>**$p=0.0277$**<br>$p=0.0975$ | T10: WT vs LKB1<sup>fl/fl</sup><br>T10: WT vs Na<sub>v</sub>1.8<sup>LKB1</sup><br>T10: LKB1<sup>fl/fl</sup> vs Na<sub>v</sub>1.8<sup>LKB1</sup><br>T11: WT vs LKB1<sup>fl/fl</sup><br>T11: WT vs Na<sub>v</sub>1.8<sup>LKB1</sup><br>T11: LKB1<sup>fl/fl</sup> vs Na<sub>v</sub>1.8<sup>LKB1</sup><br>T12: WT vs LKB1<sup>fl/fl</sup><br>**T12: WT vs Na<sub>v</sub>1.8<sup>LKB1</sup>**<br>T12: LKB1<sup>fl/fl</sup> vs Na<sub>v</sub>1.8<sup>LKB1</sup><br>T13: WT vs LKB1<sup>fl/fl</sup><br>T13: WT vs Na<sub>v</sub>1.8<sup>LKB1</sup><br>T13: LKB1<sup>fl/fl</sup> vs Na<sub>v</sub>1.8<sup>LKB1</sup> | $p=0.3430$<br>$p=0.4273$<br>$p=0.9865$<br>$p=0.4644$<br>$p=0.9754$<br>$p=0.6660$<br>$p=0.1008$<br>**$p=0.0003$**<br>$p=0.1141$<br>$p=0.9667$<br>$p=0.9136$<br>$p=0.9843$ |
|  |  |  |  | L1: WT vs LKB1<sup>fl/fl</sup> | *p*=0.7610 |
|  |  |  |  | L1: WT vs Na<sub>v</sub>1.8<sup>LKB1</sup> | *p*=0.7491 |
|  |  |  |  | L1: LKB1<sup>fl/fl</sup> vs Na<sub>v</sub>1.8<sup>LKB1</sup> | *p*=0.9971 |

We then immunostained for Na_v_1.8 as a marker for sensory neurons and tyrosine hydroxylase (TH) as a marker for sympathetic neurons and assessed the colocalization of each of these populations with retrograde fluorogold staining. WT mice had increased Na_v_1.8/fluorogold colocalization at lower DRGs (T13 and L1), with less contribution from T10-T12 DRGs, whereas LKB1^fl/fl^ mice have similar numbers of Na_v_1.8/fluorogold double-labelled neurons across all DRGs. Interestingly, Na_v_1.8/fluorogold colocalization was significantly reduced in Na_v_1.8^LKB1^ females compared to WT (T13 and L1) and LKB1^fl/fl^ (T12, T13, and L1) females (Figure 3C, Table 4). There were no differences in the number of TH/fluorogold double-labelled neurons between WT, LKB1^fl/fl^, and Na_v_1.8^LKB1^ mice, with the exception that Na_v_1.8^LKB1^ mice had significantly fewer TH/fluorogold neurons at T12 compared to WT, but not LKB1^fl/fl^ mice (Figure 3D, Table 4). These data provide novel evidence that LKB1 activity is an important contributor to ovarian innervation by peripheral sensory neurons., Thus, sensory neuron innervation from DRG neurons may be a promising future target for modulation of fertility in females.

### Histological analysis of reproductive organs in Na_v_1.8^LKB1^ males and females

Although the increased litter sizes were observed specifically when female Na_v_1.8^LKB1^ breeders were paired with LKB1^fl/fl^ males, we wanted to assess whether the removal of LKB1 from sensory neurons had any impact on overt physiology in both male and female mice. At six weeks of age, we weighed male and female Na_v_1.8^LKB1^ and LKB1^fl/fl^ mice immediately prior to euthanasia, with vaginal lavage was also performed immediately prior to euthanasia, such that females in follicular and luteal phases of the estrous cycle were represented in each group. Male and female reproductive tracts (testes and epididymis for males and uterus, fallopian tubes, and ovaries for females) were dissected and weighed immediately following their extraction. As reported previously, there were no overt differences in body weight between strain or genotype groups in either male or female mice (Supplementary Figure 2A-B, Supplementary Table 2) (Garner & Burton, 2022). Age matched WT (C57BL/6J) females had significantly lower reproductive organ weight compared to LKB1^fl/fl^ and Na_v_1.8^LKB1^ females (Supplementary Figure 2C, Supplementary Table 2). There was no such difference between LKB1^fl/fl^ and Na_v_1.8^LKB1^ females, indicating that LKB1 in peripheral sensory neurons does not affect overt uterine physiology. For male mice, there were no differences in testes and epididymis weights (Supplementary Figure 2D, Supplementary Table 2).

### Histological analysis of ovary morphology *in* Na_v_1.8^LKB1^ females

Because Na_v_1.8^LKB1^ female mice have significantly larger litters than LKB1^fl/fl^ littermate controls, we performed a histological analysis on the ovaries from these groups of mice to investigate follicular turnover, corpus luteum formation, and perigonadal adipocyte size. Measurement of ovary sizes immediately following dissection showed that Na_v_1.8^LKB1^ ovaries are significantly larger compared to LKB1^fl/fl^ ovaries (Figure 4B, Table 5). Next, we used H&E-stained ovary sections to assess follicle dynamics and the presence of corpora luteum (CL). There were no differences in the number of CL, which is a transient structure formed after ovulation that produces progesterone and supports the maintenance of pregnancy (Figure 4C, Table 5). Ovarian follicles have a well-characterized life cycle proceeding from the growth stage (primary and secondary follicles) to the maturation stage (pre-antral and antral) as they are selected for either ovulation or degeneration. We first assessed the number of follicles at each of these stages (independent of estrous cycle phase) and found that although there were no significant group differences in the number of follicles at any particular stage, Na_v_1.8^LKB1^ mice had significantly more total follicles compared to LKB1^fl/fl^ mice (Figure 4E, Table 5). This indicates that sensory neurons may play a role in regulating follicular turnover, without necessarily having a direct role in rates of ovulation.

**Figure 4.**
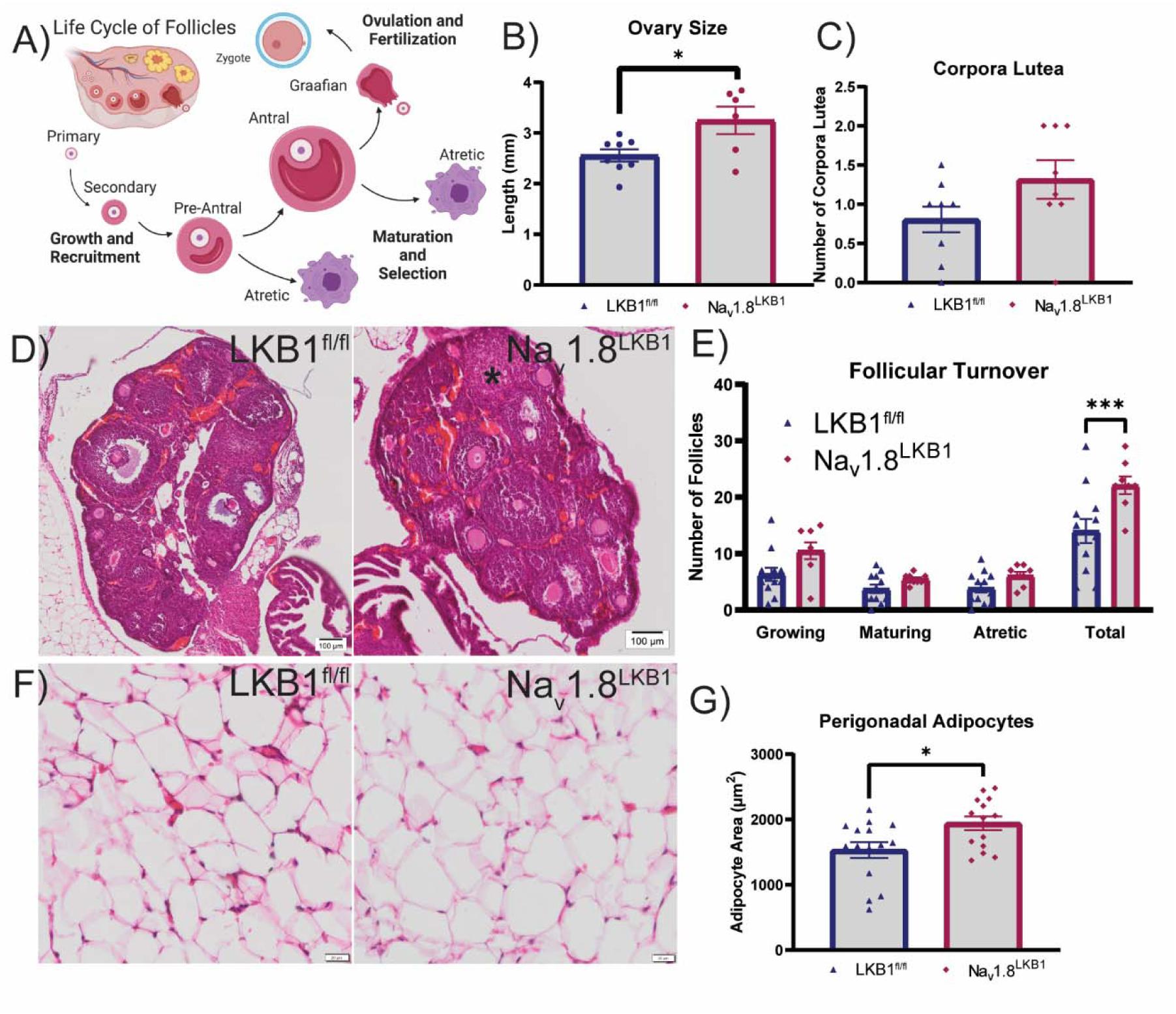
Histological analysis of ovary morphology in Na_v_1.8^LKB1^ females. (A) Graphic depiction of assessment of the follicular life cycle. (B) Ovary size as measured immediately following extraction (LKB1^fl/fl^: mean ± SEM = 2.6 ± 0.12 mm, n=8 females; Na_v_1.8^LKB1^: mean ± SEM = 3.2 ± 0.27 mm, n=6 females). (C) Number of corpora lutea (CL) per ovary (LKB1^fl/fl^: mean ± SEM = 2.0 ± 0.45 CL, n=6 females; Na_v_1.8^LKB1^: mean ± SEM = 2.1 ± 0.55 CL, n=8 females). (D) Representative images of H&E-stained ovary sections. Scale bar = 100 µm. *=CL. (E) Number of follicles at each stage. (LKB1^fl/fl^ n=12 females; Na_v_1.8^LKB1^ n=8 females) For each group, n=5-10 technical replicates/sections. (F) Representative images of H&E-stained perigonadal adipocytes. Scale bar = 20 µm. (G) Size (area in µm^2^) of perigonadal adipocytes. (LKB1^fl/fl^: mean ± SEM = 1531 ± 121.8 µm^2^, n=14 females; Na_v_1.8^LKB1^: mean ± SEM = 1943 ± 104.2 µm^2^, n=15 females). For each group, n=5-10 technical replicates (sections) with minimum 60 cells analyzed per section. *p<0.05, ***p<0.0001 by unpaired two-tailed t-test (B, C, G) or ordinary two-way ANOVA (E).

**Table 5.**
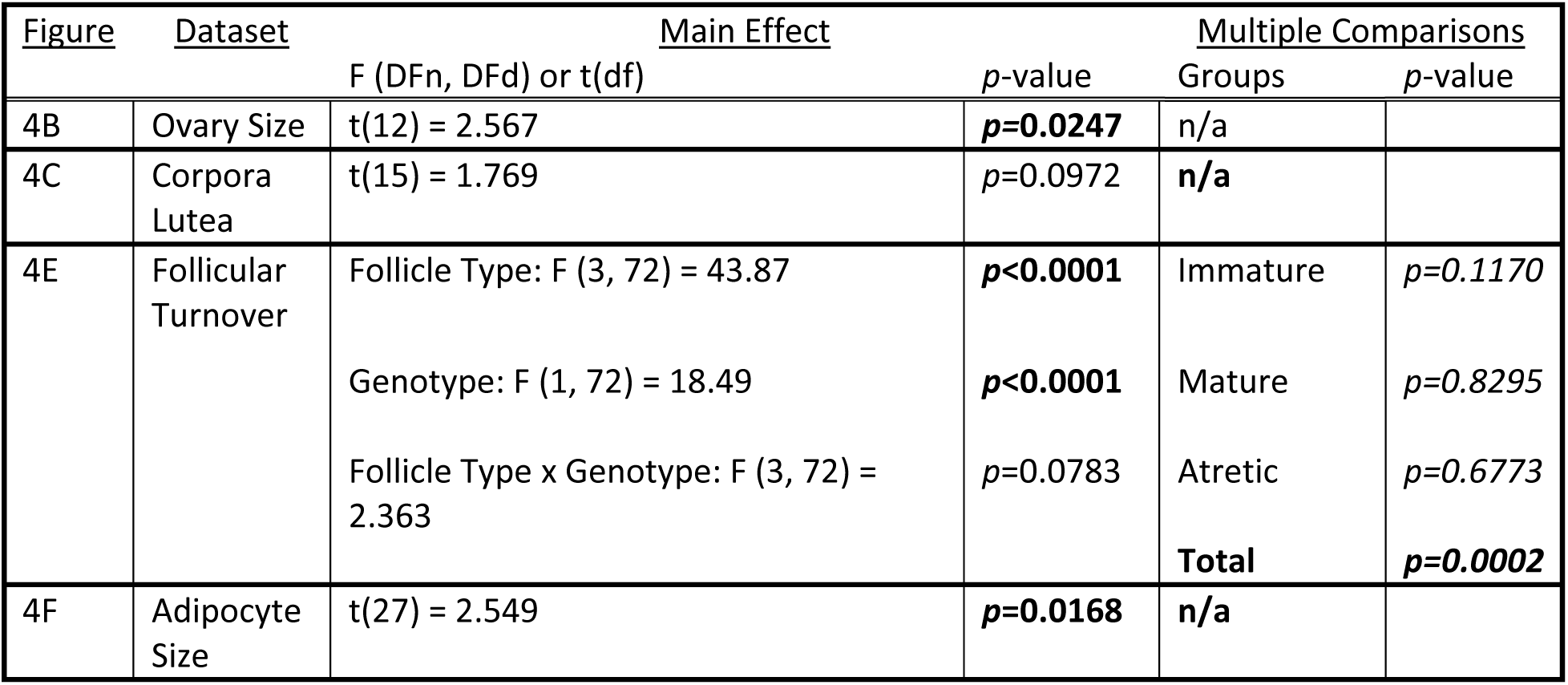
Statistics corresponding to Figure 4. Ovary size, number of corpora lutea, and adipocyte size were analyzed using an unpaired two-tailed t-test. Follicular turnover was analyzed using an ordinary two-way ANOVA with *post hoc* Tukey’s multiple comparisons. Significance was set at *p*<0.05 for all datasets. All significant results are in bold.

Disorders such as polycystic ovarian syndrome (PCOS) present with metabolic abnormalities and increased adiposity, which imply a communication between perigonadal adipose tissues and the ovaries (Agarwal *et al*., 2015; Guastella *et al*, 2010; Hohos *et al*, 2018; Lotti *et al*, 2021). Although we have previously shown that LKB1^fl/fl^ and Na_v_1.8^LKB1^ female mice have similar whole-body metabolism, we were now interested to determine if changes in perigonadal adipose would present with enhanced fertility in Na_v_1.8^LKB1^ females. We observed that Na_v_1.8^LKB1^ females had larger perigonadal adipocytes than LKB1^fl/fl^ mice, despite having comparable body weight (Figure 4G, Table 5); However, we did not assess the weight of the total fat pad. Increased adiposity is typically associated with modulated fertility; thus, the communication between perigonadal adipocytes and the ovaries in Na_v_1.8^LKB1^ females may be altered due to their reduced ovarian innervation.

### Follicular dynamics across the estrous cycle in Na_v_1.8^LKB1^ ovaries

To further examine alterations in follicular development in the Na_v_1.8^LKB1^ ovary, we analyzed the number and size of follicles in H&E-stained whole ovary sections and performed genotype comparisons of the number and size of follicle types across each phase of the estrus cycle. In Na_v_1.8^LKB1^ ovaries we found that each examined follicle type showed altered growth patterns across the estrus cycle compared to follicles from LKB1^fl/fl^ ovaries. Interestingly, Na_v_1.8^LKB1^ females had a greater abundance of primary immature follicles during diestrus versus secondary follicles estrus/metestrus (Supplementary Figure 3A and 3F, Supplementary Table 3). Primary follicles in Na_v_1.8^LKB1^ mice were also significantly larger, although secondary follicles did not differ in size (Supplementary Figure 3B-E, G-J, Supplementary Table 1). The growth of immature follicles is gonadotropin-independent and may thus be more responsive to neuronal signaling. Overall, Na_v_1.8^LKB1^ females had more pre-antral follicles than LKB1^fl/fl^ females, notably so during diestrus. There were no group differences observed in the size of pre-antral follicles (Supplementary Figure 3K-O, Supplementary Table 3), nor did we see any differences in the number or size of antral follicles between LKB1^fl/fl^ and Na_v_1.8^LKB1^ mice in any phase (Supplementary Figure 3P-T, Supplementary Table 3). We next turned our attention to atretic follicles, aiming to discern whether Na_v_1.8^LKB1^ females exhibited an altered pattern of follicular selection for ovulation. Interestingly, while Na_v_1.8^LKB1^ mice had slightly more atretic follicles during the follicular phases of the cycle, LKB1^fl/fl^ mice had significantly more atretic follicles during diestrus (Supplementary Figure 3U-Y, Supplementary Table 3). The relative reduction in atretic follicles in Na_v_1.8^LKB1^ mice during diestrus suggests that Na_v_1.8^LKB1^ females have a greater selection-to-degeneration ratio than their littermate controls, thus supporting larger litters. These data support a role for sensory innervations in regulating follicular turnover, particularly regarding the growth of immature follicles and the suppression of atresia in maturing follicles.

To assess whether the observed differences in follicle size were due to precursor oocyte size or to effects of the support cell network, we measured oocyte size in H&E-stained ovary sections. There were no group differences in the size of oocytes in primary follicles, pre-antral follicles, or antral follicles at any phase of the estrous cycle; however, oocytes from secondary follicles were smaller in Na_v_1.8^LKB1^ females (Supplementary Figure 4, Supplementary Table 4). This observation stands in direct contrast to the larger primary follicles seen in Na_v_1.8^LKB1^ females, suggesting that the increased follicular size is due to granulosa cell size and not the oocyte itself.

## Discussion

We present the novel finding that modulation of sensory neurons via removal of LKB1 directly affected breeding in female mice via changes in estrous cycle, ovarian growth, and follicular development. Removal of LKB1 from sensory neurons resulted in a decrease an increase in litter size and decrease in the breeder age for the first pregnancy. These reproductive changes were only seen in female breeders with LKB1-null sensory neurons. Thus, when male mice with LKB1-null sensory neurons were paired with wild-type (LKB1^fl/fl^) females, litter sizes were comparable to those from WT breeding pairs Our retrograde labelling studies confirmed that Na_v_1.8 neurons directly innervate the ovary from the DRG, and that this ovarian innervation declines in the absence of LKB1. These findings show a sex-specific interaction between LKB1 in sensory neurons and fertility specifically in female breeders.

The role of sensory neurons in modulating reproduction and ovarian function is not well understood. Several research groups have reported on the innervation of rodent ovaries by sensory neurons: the primary sensory contribution to the ovary is thought to be calcitonin gene-related peptide (CGRP)-containing nerve fibers (Burden *et al*., 1983; Calka *et al*., 1988; Klein & Burden, 1988). CGRP, a neuropeptide expressed by peptidergic sensory neurons, has important roles in mediating neurogenic inflammation and vasodilation, both of which are to regulation of steroidogenesis and follicle development (Hanada *et al*, 2011; Rosas *et al*., 2018; Russell *et al*, 2014). CGRP and Na_v_1.8 have up to 80% colocalization rodent sensory ganglia (Patil *et al*, 2018). In DRGs, both Na_v_1.8 and CGRP in peptidergic C-fibers are involved in the detection of inflammation and noxious stimuli (Chen *et al*, 2020; Lenert *et al*., 2021a; Pinho-Ribeiro *et al*., 2017). These neurons have direct effects on tissue physiology during disease and infection processes by detecting changes in the target tissue environment (Chiu *et al*., 2013; Garner & Burton, 2022; Huang *et al*., 2021; Szabo-Pardi *et al*, 2021). Further, activated Na_v_1.8 - expressing neurons from the nodose ganglia release the neuropeptide CGRP (Jia *et al*, 2021). Vagus and DRG Na_v_1.8 neurons may have differing effects on reproductive physiology. While DRG neurons primarily detect and respond to tissue damage and inflammation, vagal neurons are mediators of the autonomic nervous system (Kupari *et al*, 2019; Morales *et al*, 2007; Sapio *et al*, 2020; Trujillo *et al*, 2015). The presence of Na_v_1.8 neurons in both the vagal nodose ganglia and DRG suggests that sensory neurons likely play a previously unappreciated role in the regulation of reproduction and fertility (Gautron *et al*, 2011).

LKB1 pathways present a promising therapeutic for fertility disorders, particularly those that feature metabolic syndromes such as PCOS (Furat Rencber *et al*, 2018; Kocer *et al*, 2014). LKB1 has multiple downstream targets that regulate cellular metabolism and function (Alessi *et al*., 2006; Molina *et al*., 2021; Shackelford & Shaw, 2009), among which the best known is AMP-activated kinase (AMPK), a highly conserved kinase that senses and responds to changes in cellular energy levels (Herzig & Shaw, 2018; Mounier *et al*, 2013; Saikia & Joseph, 2021). During low energy states, LKB1 activates AMPK to promote ATP production by shifting the cell towards glucose metabolism and lipid oxidation (Shackelford & Shaw, 2009). Sensory neurons lacking LKB1 are unable to shift energy production pathways during metabolic stress (Garner & Burton, 2022). Mitochondria, which produce ATP via oxidative phosphorylation, determine cellular metabolic output (Herzig & Shaw, 2018; Tiosano *et al*, 2019). LKB1 is an important regulator of mitochondrial homeostasis, which is essential for axon growth; neurons lacking LKB1 have reduced axon length and branching (Asada *et al*, 2007; Barnes *et al*, 2007; Burger *et al*, 2020; Huang *et al*, 2014; Shelly *et al*, 2007; Winckler, 2007). Further, LKB1 regulates mitochondrial trafficking to axon terminals and presynaptic neurotransmitter release (Courchet *et al*, 2013; Kwon *et al*, 2016). Changes in signal transduction and neurotransmitter release may accompany the reductions in ovarian innervation observed in female mice with LKB1-null sensory neurons, further reducing communication between sensory neurons and ovaries, but further studies are needed to confirm this proposition. Likely, sensory neurons lacking LKB1 are less able to detect and communicate the metabolic status of the ovary, and thus exert attenuated regulation over follicle growth and ovulation.

The relationship between metabolism and reproduction has been intensively studied, particularly as fertility disorders such as often coincide with metabolic syndrome and other metabolic abnormalities (Colledge, 2013; Lainez & Coss, 2019; Lotti *et al*., 2021). In patients with PCOS, increased body weight and insulin insensitivity correlate with poor fertility outcomes and symptom severity (Guastella *et al*., 2010; Ruddenklau & Campbell, 2019; Witchel *et al*, 2019). Impairment of steroid hormone metabolism and anovulation are typically worsened with increased body weight (Deng *et al*., 2017). Conversely, a similar phenomenon occurs in patients with anorexia nervosa and during prolonged malnutrition (Wassif *et al*., 2011). Weight gain and loss both represent significant shifts in nutritional availability, energy stores, and metabolic demand, which tend to divert energy use away from reproduction and towards immediately essential functions (Perrigo & Bronson, 1983). In WT mice, the relationship between metabolism and reproduction is less straightforward, insofar as female mice (C57BL/6J) are resistant to diet-induced weight gain and hyperglycemia (Lainez *et al*, 2018; Lenert *et al*, 2021b). Nonetheless, metabolic changes during high-fat diet consumption manifest in altered ovarian function via shifts in estrous cycling, ovarian gene expression, and steroid hormone production, even in the absence of overt weight gain and obesity (Hohos *et al*., 2018; Lenert & Burton, 2021; Negrón & Radovick, 2020). Because sensory neurons lacking LKB1 have a reduced capability to innervate tissue and an impaired response to metabolic stress, mice with LKB1-deficient sensory neurons may have impaired ability to detect and/or respond to changes in energy demand (Garner & Burton, 2022). Our previous study in LKB1-deficient mice did not show any overt differences in weight loss and regain after a 24 hour fast; however, female Na_v_1.8^LKB1^ mice showed prolonged sensitization of peripheral sensory neurons compared to LKB1^fl/fl^ littermate controls (Garner & Burton, 2022). Those results, together with current observations that Na_v_1.8^LKB1^ females have larger perigonadal adipocytes and enhanced fertility, provide further evidence that sensory neurons play an essential role in metabolic stress responses. Interestingly, a recent landmark study demonstrated that ablation of sensory neurons that innervate visceral adipose resulted in increased fat mass and an upregulation of lipogenesis-promoting genes (Wang *et al*., 2022). While those findings were in male mice, there is general concurrence with present results. In particular, Wang et al. also observed that changes to sensory innervation (via ablation) did not result in overtly altered body weight, much as we observed in male and female Na_v_1.8^LKB1^ mice. The sex-specific findings in our study highlight the need for additional research into the mechanisms whereby females regulate and respond to metabolic stress, as further understanding of these processes may pave the way for therapeutic developments for treating female fertility.

Interactions between LKB1 and sex hormones, particularly 17β-estradiol (E2), and their receptors may underly the present findings in female mice - and in our previous work. While not yet reported in neurons, several studies have observed a link between LKB1 activity and estrogen receptor (ER) signaling in other cell types. ERs (subtypes alpha, α, and beta, β) are nuclear hormone receptors expressed in reproductive (i.e., ovary, uterus, testes, epididymis) and non-reproductive (i.e., bone, immune cells, adipose tissue, liver, nervous system) tissues in both males and females (Dahlman-Wright *et al*, 2006). ERα and ERβ have markedly differing physiological effects: ERα promotes cellular proliferation and prevents weight gain and insulin resistance, while ERβ has anti-inflammatory effects and often antagonizes ERα responses (Arnal *et al*, 2017; Leitman *et al*, 2010; Paterni *et al*, 2014). Of the two subtypes, ERα has the best-established interaction with LKB1 (Chen *et al*, 2016). As a transcription factor, ERα repressed the expression of LKB1 in ERα-positive breast cancer cells, which was rescued by E2 treatment (Linher-Melville *et al*, 2012; Lipovka *et al*, 2015). LKB1 also directly modulates ERα signaling by enhancing ERα-mediated responses to E2 (Nath-Sain & Marignani, 2009). On the other hand, there is some evidence that ERβ can interact with LKB1 by modulating the activation of AMPK in response to E2 (Yang *et al*, 2019). Subtype-specific expression of ERs has been observed in DRG neurons, with ERα expression predominating in small to medium diameter DRG neurons and ERβ expression in small subpopulations of small, medium, and large diameter DRG neurons (Papka *et al*, 2001). In our study, we observed significantly reduced diestrus serum estradiol levels during in Na_v_1.8^LKB1^ female mice; however, levels in other phases of the cycle were similar to wild-type female and male mice. While sensory neurons are not directly responsible for steroid hormone production, the lack of LKB1 in these cells likely modulates their response to circulating hormones such as E2.

While we believe E2 has the greatest potential among hormones for interacting with LKB1 in sensory neurons, we do not discount possible effects of LKB1 on other steroid hormones. LKB1-knockdown enhances androgen synthesis in theca cells, which are ovarian follicular endocrine cells (Magoffin, 2005). Interestingly, these theca cells preferentially express the ERα subtype (Dahlman-Wright *et al*., 2006). In granulosa cells, LKB1-knockdown reduced aromatase activity, thereby diminishing the conversion of testosterone to estradiol: aromatase expression is often inversely related to LKB1 activity (Brown *et al*, 2009; Ham *et al*, 2017). In adipocytes, treatment with the androgens testosterone and dihydrotestosterone (DHT) inhibited LKB1 expression, thereby reducing activation of AMPK (McInnes *et al*, 2012).

A limitation of our study is that we made our assessments of ovarian innervation and histology in non-pregnant mice. Our goal was to focus on the relationship between ovarian physiology and sensory neurons in relation to female fertility; however, we acknowledge that there is much to be gained from investigating the role of sensory neurons in the establishment and maintenance of pregnancy. The uterus also has sensory innervation and likely receives a contribution from DRG neurons (Chaban *et al*, 2007; Dodds *et al*, 2021; Herweijer *et al*, 2014; Zoubina & Smith, 2001). We would presume reduced uterine sensory neuron innervation would likely favor healthy pregnancies, as increased sensory innervation of the uterus occurs in reproductive disorders such as endometriosis (Arnold *et al*., 2012; Zhang *et al*, 2010). A limitation in studying uterine physiology in this context is that the rodent uterus lacks the cyclical shedding of the endometrium and resulting menstruation seen in humans and non-human primates; thus, the role sensory of neurons in preparing for and maintaining a healthy pregnancy may differ in rodents and primates (Lim & Wang, 2010). Interestingly, cyclic changes in the density of reproductive tract innervation have been observed across the reproductive cycle and during pregnancy in both rodent and humans (Mónica Brauer & Smith, 2015; Tingåker & Irestedt, 2010). Thus, there remains much to be established regarding the dynamics of sensory feedback in relation to fertility.

In conclusion, modulating the metabolic profile of sensory neurons via deletion of LKB1 resulted in a phenotype of enhanced fertility in female mice, increased litter sizes, and younger pregnancies. Breeding a male with LKB1-null sensory neurons with a WT female did not alter fertility and produced similar litter sizes to wild-type breeders (Figure 1A). Thus, this sex-specific reproductive phenotype highlights a novel role for sensory neurons in modulating fertility. Na_v_1.8-expressing sensory neurons innervate the ovary from the DRG, but this innervation is reduced in the absence of LKB1. These data support existing literature showing that LKB1 is essential for proper axonal growth and innervation, which we now extent to the case of peripheral sensory neurons. This LKB1 expression in sensory neurons present a promising target for understanding neuronal regulation of female fertility.

## Materials and Methods

### Animals

Unmated female mice (6-10 weeks old) of two different strains were used for all experiments, except for breeding studies (see below). WT C57BL/6J mice were purchased from Jackson Laboratories (stock no. 000664) and used to establish our in-house breeding colony at UT-Dallas. Transgenic mice (C57BL/6) expressing NLS-Cre recombinase under control of the *Scn10a* (Na_v_1.8) promoter were obtained initially from Professor John Wood (University College London) but are commercially available from (Ifrafrontier, EMMA ID: 04582). Although most Na_v_1.8-expressing neurons are nociceptors (C-fibers), Na_v_1.8 is expressed in small diameter neurons in the dorsal root ganglia, trigeminal ganglia, and nodose ganglia and can therefore be broadly identified as peripheral sensory neurons (Gautron *et al*., 2011; Patil *et al*., 2018; Stirling *et al*, 2005). Previous characterization of these mice demonstrated normal electrophysiological properties and expression of Cre recombinase in approximately 75% of Na_v_1.8-positive neurons in the DRG (Stirling *et al*., 2005; Szabo-Pardi *et al*., 2021). Genetically modified LKB1 flox mice (FVB;129S6-*Stk11^tm1Rdp^*/Nci; LKB1^fl/fl^), which have loxP sites flanking exons 3-6 of the *Stk11* gene, were purchased from the Frederick National Laboratory for Cancer Research (Strain No. 01XN2). LKB1^fl/fl^ mice were bred with Na_v_1.8cre heterozygous mice to produce cell-specific LKB1 knockouts (Na_v_1.8^LKB1^) (Bardeesy *et al*, 2002; Garner & Burton, 2022). Our breeding strategy was such that Na_v_1.8^LKB1^ animals were homozygous for the LKB1 floxed gene (LKB1^fl/fl^) and expressed one copy of Na_v_1.8cre. Littermate controls (LKB1^fl/fl^) expressed two copies of the LKB1 floxed gene and had normal LKB1 expression in the absence of Na_v_1.8cre (Garner & Burton, 2022). WT mixed strain mice (C57BL/6J x FVB:129S) were bred in-house and used as a strain control for fertility assessments. ICR (Envigo, Indianapolis, IN) breeders served as an outbred strain control. Fertility metrics from ICR breeders were generously provided by Theodore Price at the University of Texas at Dallas.

All animals used in this study were bred in-house and group housed (3-5 per cage) in polypropylene cages. Animal housing rooms were maintained at a temperature of 21 ± 2°C under a 12h light-dark cycle (lights on at 6 AM and off at 6 PM). Mice had *ad libitum* access to water and standard rodent chow (LabDiet ProLab RMH 1800). All procedures used in this study were performed in accordance with the National Institutes of Health Guidelines for the Care and Use of Laboratory Animals and in accordance with ARRIVE guidelines (National Research Council Committee for the Update of the Guide for the & Use of Laboratory, 2011; Percie du Sert *et al*, 2020). All procedures used in this study were approved by the University of Texas at Dallas Institutional Animal Care and Use Committee protocols 16-07 (breeding) and 17-10 (experiment).

### Polymerase chain reaction (PCR)

Animals were weaned between 21-28 days of age, at which time a tail clip biopsy (∼1 mm) was collected. DNA was extracted from tail clips by suspension in 75 µl extraction buffer (25 mM NaOH, 0.2 mM EDTA in ddH_2_O) with incubation at 95°C for one hour. An equal amount of neutralization buffer (40 mM Tris-HCl, pH 5.5) was added following incubation. DNA samples were stored at −20 °C until genotyping. PCR was performed using JumpStart REDTaq Ready Mix (Sigma, Cat#P0982-800RXN) and the primers listed below. Genotyping for Na_v_1.8cre was performed as a single reaction, whereas LKB1 LoxP and WT reactions were performed separately. The LKB1-flox gene was amplified using the primer pair LKB1rdp_COM 5’-GAG ATG GGT ACC AGG AGT TGG GGC T −3’ and LKB1rdp_WT 5’- GGG CTT CCA CCT GGT GCC AGC CTG T - 3’. The LKB1 wild-type gene was amplified using the primer pair LKB1rdp_COM 5’- GAG ATG GGT ACC AGG AGT TGG GGC T −3’ and LKB1rdp_MUT 5’- TCT AAC GCG CTC ATC GTC ATC CTC GGC −3’. The Na_v_1.8cre gene was amplified using the custom primer pair newNAV-WT-Fwd 5’ GAT GGA CTG CAG AGG ATG GA −3’ and newNAV-Cre-Rev 5’- CGT ATA TCC TGG CAG CGA TC −3’. The Na_v_1.8 WT gene was amplified using the custom primer pair newNAV-WT-Fwd 5’ GAT GGA CTG CAG AGG ATG GA −3’ and newNAV-WT-Rev 5’- GGT GTG TGC TGT AGA AAG −3’. Samples were run on a 2% agarose (VWR, Cat# MPN605-500G) and 1X Tris-Acetate-EDTA (TAE) gel and imaged using a ChemiDoc apparatus (BioRad) and ImageLab. DNA Ladder (100 bp, Thermo Scientific GeneRuler 100 bp) was used for size determination. The Na_v_1.8cre and Na_v_1.8 WT bands were at 800 and 500 bp, respectively (Supplemental Figure 1A-B), whereas the LKB1 mutant and LKB1 WT bands were at 300 and 220 bp, respectively. In the LKB1 WT reaction, a band at 600 bp appeared in the presence of LoxP sites (Supplemental Figure 1B) (Bardeesy *et al*., 2002).

### Breeding History

Breeding pairs were initiated when animals were between 5-6 weeks of age, except for a small cohort (analyzed separately) that was initiated at 20 weeks of age. Breeding pairs outside of these age ranges were excluded from analyses. All breeding pairs are set up as trios, with one male paired with two females, to promote pair nursing, cross fostering, and pup health. Per facility and husbandry guidelines, litters older than ten days were separated with their respective dam in cases of overcrowding. Our breeding strategy was such that Na_v_1.8^LKB1^ males were paired with LKB1^fl/fl^ females or LKB1^fl/fl^ males with Na_v_1.8^LKB1^ females (Figure 1A). WT breeding pairs (C57BL6/J, mixed strain, and ICR) had both male and female WT breeders. At no point were male breeders removed from the breeding cages.

Breeders were carefully tracked by experimenters to ensure accuracy of breeding data. Breeders were handled only by experimenters, with glove changes between cages to reduce animal stress by minimizing scent transfer. Breeding cages were cleaned exclusively by familiar experimenters. Per facility and husbandry guidelines, all cages were provided with standard rodent bedding, nesting material for enrichment, and a red igloo nesting box. When initiating breeding pairs, mice were placed in a clean cage with fresh food, bedding, and nesting material. Female estrous cycles were not tracked prior to the initiation of breeding pairs.

All breeding information was collected for individual female breeders: data was never averaged for two females in the same trio. All genotypes referenced in breeding data refer to the genotype of the female breeder. The age at first litter was defined as the age of the female breeder on the date of birth of her first viable litter, i.e., precocious puberty, as noted in Na_v_1.8^LKB1^ females, was marked in cases where female weanlings were impregnated prior to weaning (<4 weeks old). Conversely, days to first litter was defined as the number of days between breeding initiation to the birth of the first viable litter per female in each breeding pair. Litter sizes were determined by counting the number of viable pups born from each female and were tracked for the first five litters for each female. Rare occurrences (<1%) of non-viable pups (death due to cannibalism, maternal neglect, etc.) were not counted for this study. Litter frequency was measured as the number of days between consecutive viable litters for each female breeder. Breeding schematics can be found in **Table 1** and **Supplementary Table 1**.

### Estrous Cycling

Estrous cycling was assessed over a ten-day period via cytology of vaginal lavage samples. Starting at six weeks of age in non-mated female mice, samples were collected twice per day between 8-10 AM and 5-7 PM for ten consecutive days. Vaginal lavage samples were collected using previously described methods by experimenters blinded to genotype (Byers *et al*, 2012; Cora *et al*, 2015; Lenert *et al*., 2021b). To summarize, vaginal lavage with 20 µL of sterile saline was collected onto charged microscope slides (Fisher, Cat#12-544-7) and allowed to air dry overnight. Samples were stained with toluidine blue for visualization of cells and imaged using brightfield microscopy. Classification of cycle phase was performed by experimenters blinded to genotype. Phases were determined by cell type and density present in the stained vaginal lavage samples. Proestrus smears consist primarily of nucleated epithelial cells, whereas estrus smears consist primarily of anucleated or “cornified” epithelial cells that appear in high density and often in clumps. Metestrus smears consist of a mixture of anucleated epithelial cells and neutrophils, with few little nucleated epithelial cells, and are usually of high cell density. Diestrus smears consist primarily of neutrophils and range from moderate to low cell density. Transitory phases were marked as such. For example, a smear with an equal number of nucleated and anucleated epithelial cells would be classified as a proestrus-estrus transition and denoted “P/E”. Complete cycle tracking information can be found in **Table 6**. To compare estrous cycle tracking between groups, each phase is represented as a percentage, where the number of samples in each phase was divided by the total number of collected samples per mouse. For example, if two of 20 samples over the ten-day period were in proestrus, this would be represented as 20%. Cycle length was measured as the number of days between consecutive proestrus smears.

**Table 6.**
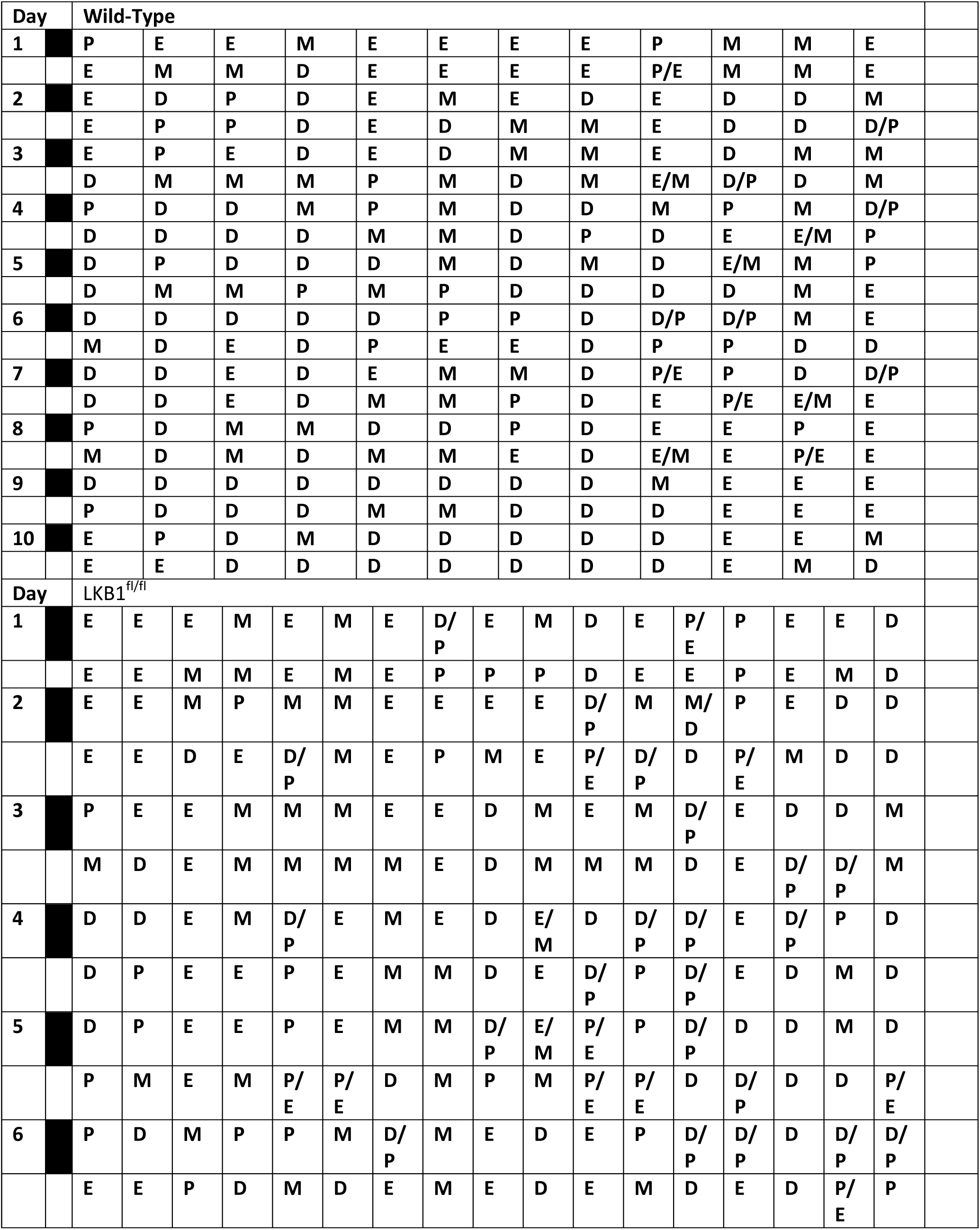
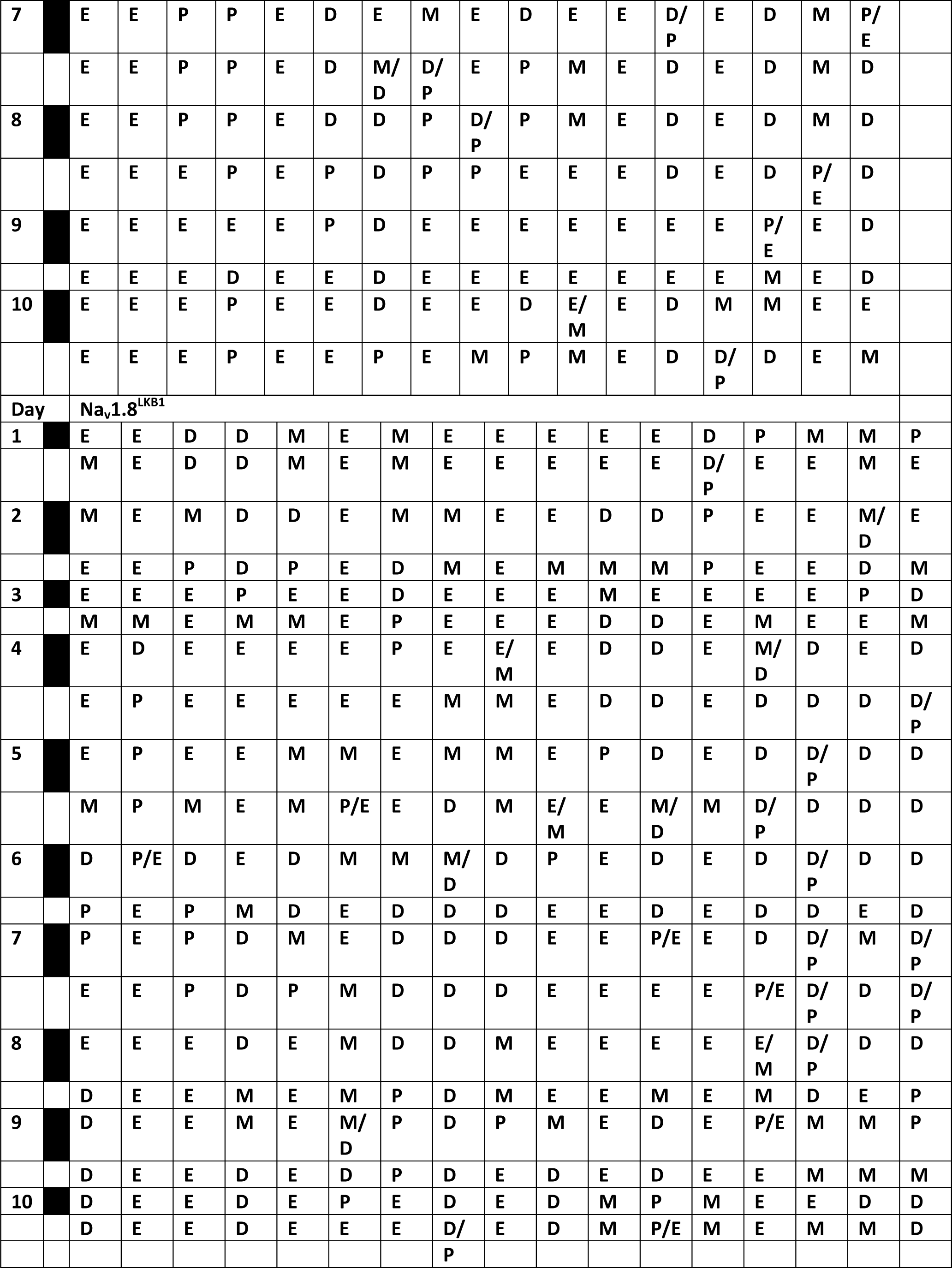
Complete estrous cycle tracking data. P=Proestrus, E=estrus, M=metestrus, D=Diestrus. Shaded cells indicate samples collected during the dark cycle period. Each column represents one female mouse.

### Estradiol ELISA

Serum was isolated from tail blood collected in the morning (9 AM-11 AM) immediately following vaginal lavage collection. Tail blood was collected using microvette EDTA-coated tail vein capsules (Sarstedt, Cat#16.444.100). Serum was isolated via centrifugation at 14,000 RPM for 15 minutes at 4 °C. Serum samples were immediately stored at −80 °C. Quantification of serum estradiol was performed using the Crystal Chem Estradiol ELISA kit (Cat#80548) according to the manufacturer’s instructions. The assay has a detectability range of 5-1280 pg/mL and a CV of 10%. The samples were assayed at a 1:3 dilution in 0 pg/mL standard buffer.

### Retrograde Tracing

To determine the extent of sensory innervation to the ovary, we performed retrograde tracing studies on WT, LKB1^fl/fl^, and Na_v_1.8^LKB1^ females. Fluorogold (Hydroxystilbamidine, Biotium, Cat#80014) was diluted in ddH2O to a 0.4% solution and stored at 4 °C in opaque/black tubes (Black Beauty Microcentrifuge Tubes, Argos Technologies, 1.5mL, Cat#47751-688) (dos Santos *et al*, 2022; Miller *et al*, 2021; Mondello *et al*, 2016; Schmued & Fallon, 1986). All surgical tools were autoclaved before use. During the surgery, tools were sterilized using a Dry Glass Bead Sterilizer (Stoelting, Cat#50287) and 100% anhydrous ethanol (Decon Laboratories, CAS#64-17-5, Cat#2701). The mice were heavily sedated using isoflurane (5% induction, 3% maintenance). The surgical site was shaved and cleaned with 10% povidone iodine prep solution (USP equivalent to 1% available iodine, Dynarex, Cat#1416) and 100% anhydrous Ethanol. Via laparotomy, the skin and peritoneum were incised with a #10 scalpel blade (Sigma-Aldrich, Cat#S2646) and the ovary and perigonadal fat pad were exposed. A 5 µL volume of the 0.4% Fluorogold solution was injected into the left ovary using a 30-gauge needle mounted on a 25 µL Hamilton syringe. The peritoneum was sutured closed using a 5-0 silk suture (VWR, MV-682) and the skin was then closed using an AutoClip Kit with 9 mm clips (Fine Science Tools, Cat#12020-00). Mice were given a subcutaneous injection of Gentamicin (10 mg/kg, sterile-filtered, Sigma-Aldrich, Cat#G1272-100mL) as a preventative antibiotic into the nape of the neck with a 25-gauge needle using a 1 mL syringe and returned to their home cages to recover. Mice were also given 0.125 mg Meloxicam (Bio-Serv, Cat#M275-050D) for post-surgical pain and monitored daily for infection or malaise.

### Immunohistochemistry

For retrograde tracing experiments, ten days following Fluorogold injection, mice were euthanized following the University of Texas at Dallas IACUC procedure. Bilateral DRGs (T7-L5) were collected and post-fixed in 4% paraformaldehyde (PFA) for 4-6 hours. DRGs were cryoprotected in 30% sucrose (in 1X phosphate-buffered saline (PBS)) for 48 hours and frozen in OCT. Frozen DRGs were serially sectioned on a cryostat at 14 µm and mounted onto charged microscope slides. Each slide contained ipsilateral and contralateral DRG from the same level (i.e., all L1 samples) from each group. Frozen DRG sections were stored at −20 °C prior to immunohistochemistry. All samples were sectioned such that every fifth section was represented on one slide, spanning the entire DRG. Frozen sections were dried out for 15 minutes at room temperature (RT) prior to rehydration of the samples with 1X PBS. For antigen retrieval, rehydrated sections were incubated in heated 10 mM sodium citrate buffer (in 1X PBS, pH 6.0; Sigma, Cat#C8532-500G) for 5 minutes, three times in succession. Sections were then incubated in blocking buffer (3% normal goat serum (Gibco, Cat#16210-072), 2% bovine serum albumin (VWR, Cat#97061-416), 0.1% Triton X-100 (Sigma, Cat#X100), 0.05% Tween-20 (Sigma-Aldrich, Cat#P1379), and 0.1% sodium azide (Sigma, Cat#RTC000068) in 1X PBS, pH 7.4) for two hours at RT followed by overnight incubation at 4 °C in a primary antibody solution consisting of Anti-Na_v_1.8, Anti-tyrosine hydroxylase (TH), and Anti-Fluorogold diluted in blocking buffer. Sections were then washed three times in 1X PBS + 0.05% Tween-20 (PBS-T) and incubated in secondary antibody solution diluted in blocking buffer for two hours at RT, protected from light. Sections were then washed three times in 1X PBS-T, incubated in DAPI (1:5000) for one minute, washed once with 1X PBS, and mounted with Gelvatol. Mounted sections were stored at RT overnight in dark and then stored at 4 °C until imaging. Details about antibodies can be found in **Table 7**.

**Table 7.**
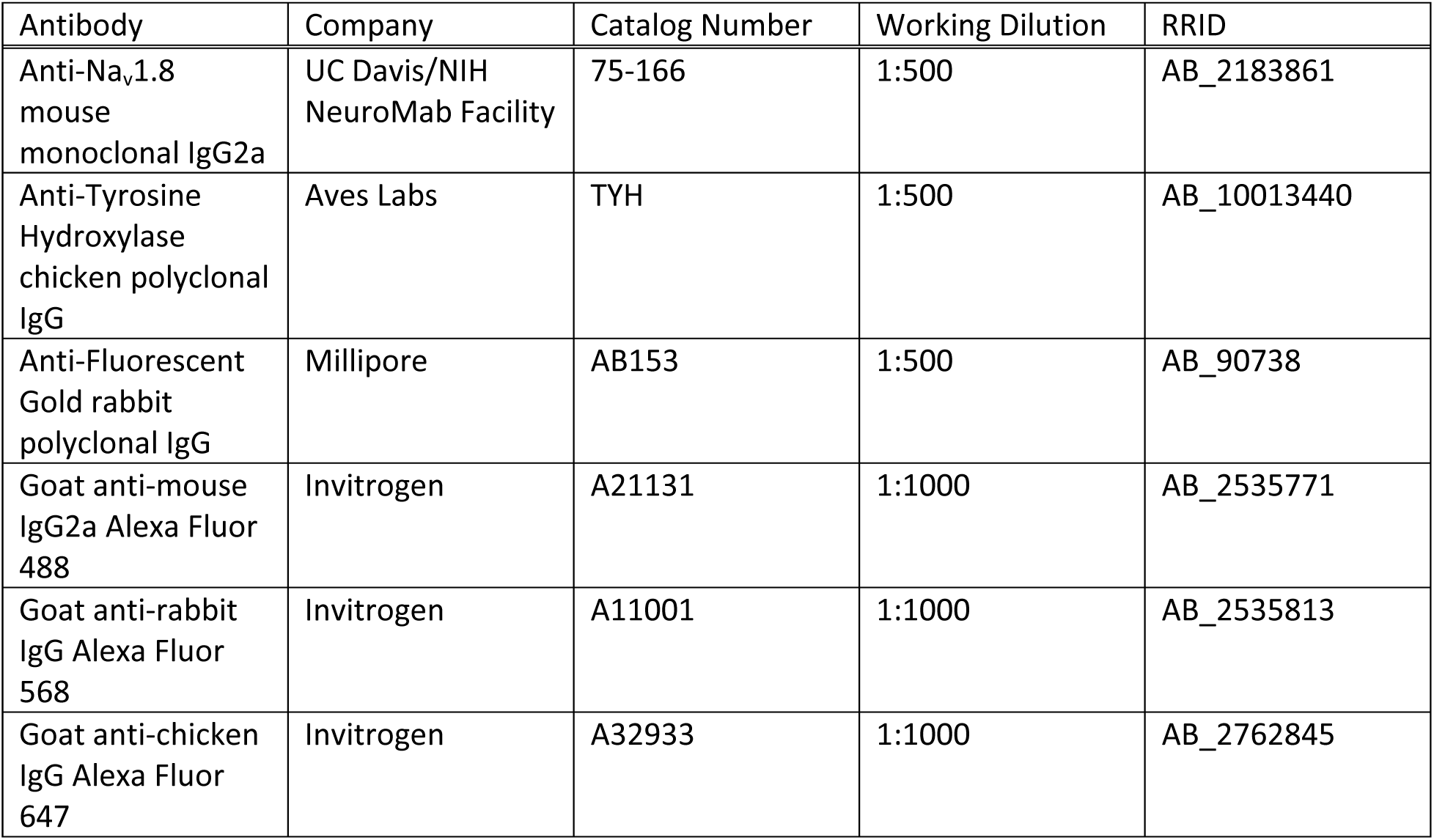
Antibodies used for immunohistochemistry.

### Image acquisition and analysis for fluorescent microscopy

Images for analysis were acquired using the Zeiss Axio Observer Microscope equipped with an Axiocam 503 B/W monochrome camera (Carl Zeiss, Inc.). Epi-fluorescent z-stack images were taken with 20x objective (NA 0.8) 0.227µm/pixel using ZEN2.5 Pro software (Carl Zeiss, Inc.). Further information regarding equipment and settings can be found in **Supplementary Table 5.** Image analysis was performed on five serial 14 µm sections per DRG using the Olympus CellSens Software using the “Count and Measure” feature. Within each subpopulation, only cells with visible nuclei DAPI staining were measured to avoid overcounting duplicate cells. First, the total number of neurons with fluorogold signal were counted. Then, we counted the fluorogold-positive neurons that colocalized with either Na_v_1.8^+^ or TH^+^ immunostaining. Experimenters were blinded to genotype for all image acquisition and analysis. Representative images for publication were acquired using the Olympus FV300RS Confocal Laser Scanning Microscope using the 30X objective (UPLSAPO30x; UPLSAPO N 30X SI OIL, NA 1.05, WD 0.8 mm w/ correction collar) using the 3D Z-stack mode on the Complete Fluoview Acquisition Software.

### Tissue Collection and Processing

For histology experiments, estrous cycle phase was determined using the measures described above immediately prior to tissue collection. For tissue collection, mice were euthanized following the University of Texas at Dallas IACUC procedure and the ovaries and surrounding perigonadal fat were collected and post-fixed in 4% PFA (made in 1X PBS) for 4-6 hours. Ovary size (both ovaries) was measured immediately following extraction using a caliper by an experimenter blinded to genotype. One ovary per mouse together with its perigonadal fat pad was embedded in paraffin and cut into 10 µm-thick sections, which were mounted on charged microscope slides. For histological analysis of ovarian and adipocyte structure, paraffin embedded sections were stained with Hematoxylin (Sigma, Cat#HHS16) and Eosin Y Solution (Sigma-Aldrich, Cat#318906) (H&E).

### Image Acquisition and Analysis of Ovaries and Adipocytes

H&E-stained tissue sections were imaged via brightfield microscopy using the Olympus VS120 Virtual Slide Microscope equipped with the 40x objective (NA 0.95, 0.33um/pixel). Analysis of 5-10 serial sections per ovary was performed using the Olympus cellSens Software. Sections were analyzed by experimenters blinded to estrous cycle phase and genotype. Follicles were classified based on a modified scheme originally proposed by Pedersen & Peters, 1986 (Gaytan *et al*, 2017; Pedersen & Peters, 1968; Westwood, 2008). In brief, follicles were classified into one of five types: primary, secondary, pre-antral, antral, or atretic (degenerating). Oocytes with a single or multiple surrounding layers of granulosa cells were classified as primary or secondary follicles, respectively. Oocytes surrounded by a defined zona granulosa and theca without the formation of a follicular antrum were classified as pre-antral follicles, whereas those with a formed follicular antrum were classified as antral follicles. Atretic follicles were determined by the presence of apoptotic cells. Adipocytes were quantified by measuring the area of at least 60 randomly selected adipocytes per section in the perigonadal fat pad.

### Statistics

All statistical analysis was performed using GraphPad Prism 9.4.0 software. All data are expressed as mean ± SEM. Statistical significance was set at *p* < 0.05 for all analysis. One-way ANOVA with *post hoc* Tukey’s multiple comparisons was used to analyze datasets for breeding metrics except for litter sizes over time, which was analyzed using Ordinary Two-Way ANOVA with *post hoc* Tukey’s multiple comparisons. Estrous cycling data (each phase analyzed separately) were analyzed using one-way ANOVA with *post hoc* Tukey’s multiple comparisons. One animal from the Na_v_1.8^LKB1^ group was removed from the estrus cycle tracking data as it was determined to be an outlier via the Grubb’s test with α = 0.05. Serum estradiol data were analyzed using two-way ANOVA with *post hoc* Tukey’s correction for multiple comparisons. Immunohistochemistry for retrograde tracing experiments were analyzed using ordinary two-way ANOVA with *post hoc* Tukey’s correction for multiple comparisons. Histological datasets were analyzed using a two-tailed unpaired t-test. Follicle and oocyte histology datasets were analyzed using ordinary two-way ANOVA with *post hoc* Sidak’s correction for multiple comparisons. Histograms for follicle histology were made by calculating the frequency distribution for each data set and plotting the percentage of samples within each bin.

## Acknowledgements

The authors thank all current and past members of the Neuroimmunology and Behavior Laboratory and Center for Advanced Pain Studies at the University of Texas at Dallas. In particular, the authors express gratitude to Natalia L dos Santos for her invaluable expertise and training in estrous cycle tracking and classification. The authors also thank Michaela Chaparro for her technical expertise and assistance. Finally, the authors thank Theodore J. Price and the Pain Neurobiology Research Group for their generous contributions to the ICR fertility metrics. This work was supported by NIH grant F99NS129173 (MEL), Eugene McDermott Graduate Fellowship 202205 (MEL), R21DK130015-01A1 (MDB), Rita Foundation Award in Pain (MDB), and the University of Texas Rising STARS program research support grant (MDB).

## Author Contributions

MEL and MDB conceptualized the study and designed the experiments. MEL acquired and analyzed data, drafted, and edited the manuscript, drafted figures, and formal analysis/data interpretation. MDB participated and supervised in all aspects of the study from conception, design, acquisition, interpretation, and manuscript preparation. All authors reviewed and approved the final manuscript.

## CRediT authorship contribution statement

Melissa E Lenert: Conceptualization, Data curation, Formal analysis, Investigation, Methodology, Writing. Michael D Burton: Conceptualization, Data curation, Formal analysis, Funding acquisition, Methodology, Resources, Supervision, Writing

## Data availability

This study included no data deposited in external repositories.

---

**Supplemental Figure 1.**
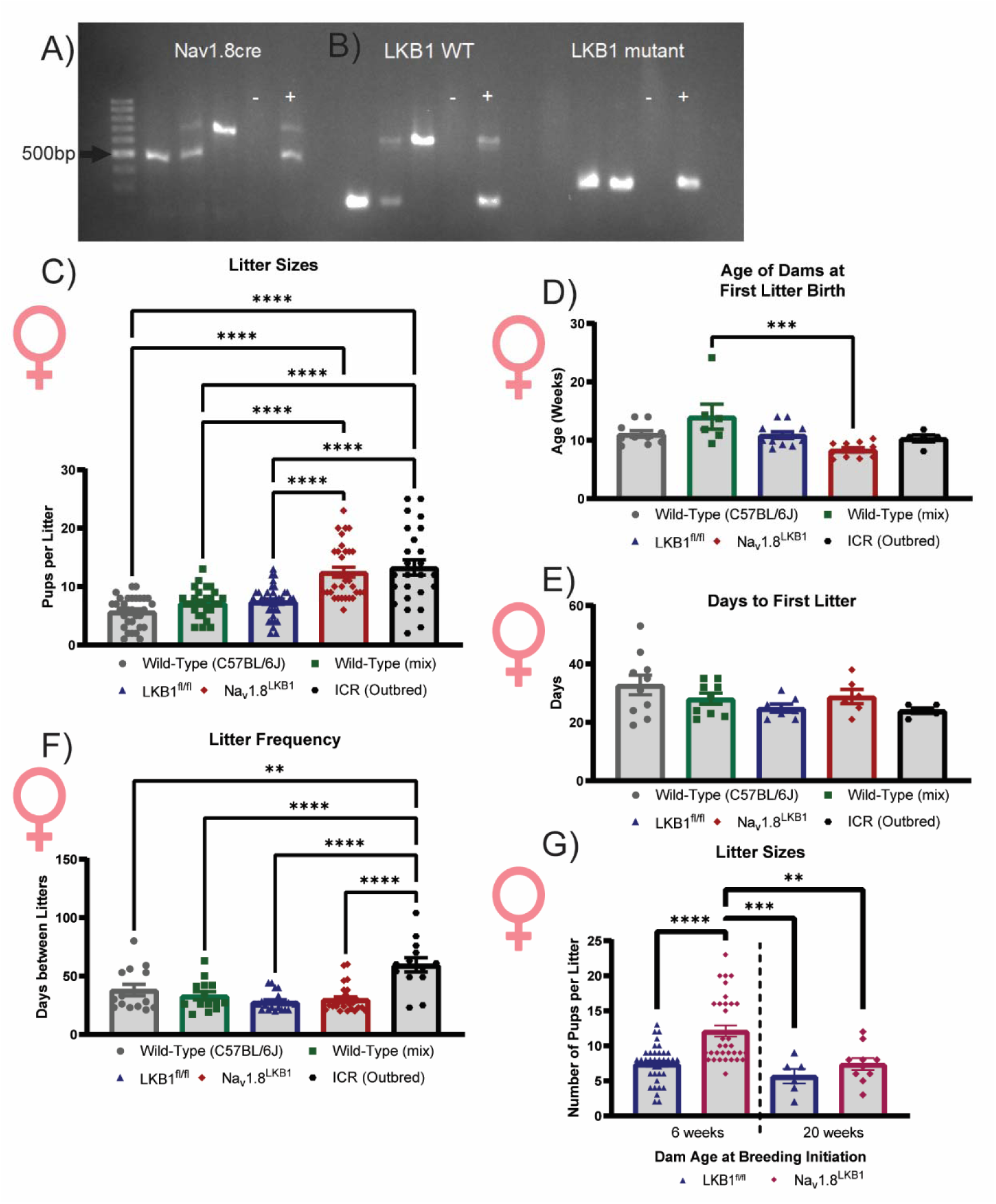
Female mice with LKB1-null sensory neurons have enhanced fertility. (A-B) Representative PCR for genotyping of Na_v_1.8cre and LKB1-loxP. (A) Na_v_1.8cre and Na_v_1.8 WT bands are at 800 and 500 bp, respectively. (B) LKB1 WT and LKB1 loxP bands at 200and 300 bp, respectively. A band at 600 bp appears in the WT reaction in the presence of loxP sites. (C) Number of pups per litter (WT: mean ± SEM = 5.7 ± 0.47 viable pups, n=31 litters from 10 females with 2-5 litters each; LKB1^fl/fl^: mean ± SEM = 7.4 ± 0.45 viable pups, n=35 litters from 11 females with 2-5 litters each; Na_v_1.8^LKB1^: mean ± SEM = 12.1 ± 0.80 viable pups, n=31 litters from 11 females with 2-5 litters each; WT mixed: mean ± SEM = 7.2 ± 0.48 viable pups, n=27 litters from 6 females with 4-5 litters each; ICR/Outbred: mean ± SEM = 13.3 ± 1.32 viable pups, n=25 litters from 5 females with 5 litters each. (D) Dam age in weeks at the birth of their first litter (WT C57: mean ± SEM = 11.1 ± 5.6 weeks, n=10 females; WT (mixed): mean ± SEM = 14.1 ± 2.14 weeks, n=6 females; LKB1^fl/fl^: mean ± SEM = 10.9 ± 0.57 weeks, n=11 females; Na_v_1.8^LKB1^: mean ± SEM = 8.4 ± 0.38 weeks, n=11 females; ICR/Outbred: mean ± SEM = 10.3 ± 0.60 weeks, n=5 females). (E) The number of days between breeding initiation and the first litter birth (WT C57: mean ± SEM = 32.8 ± 3.4 days, n = 10 females; WT mixed: mean ± SEM = 28.8 ± 2.5, n=6 females; LKB1^fl/fl^: mean ± SEM = 28.1 ± 1.9 days, n=9 females; Na_v_1.8^LKB1^: mean ± SEM = 24.9 ± 1.4 days, n=7 females; ICR/Outbred: mean ± SEM = 24.0 ± 0.95). (F) The number of days between consecutive litters per dam. (WT C57: mean ± SEM = 38.0 ± 4.96 days, n=14 from 10 females with 2-5 litters each; WT mixed: mean ± SEM = 33.1 ± 3.5 days, n=14 from 6 females with 2-5 litters each; LKB1^fl/fl^: mean ± SEM = 27.9 ± 1.7 days, n=21 from 11 females with 2-5 litters each; Na_v_1.8^LKB1^: mean ± SEM = 30.04 ± 2.24, n=26 from 11 females with 2-5 litters each; ICR/outbred: mean ± SEM = 59.6 ± 6.1 days, n=13 from 5 females with 2-3 litters each). (G) Number of litters separated by dam age at breeding initiation. (LKB1^fl/fl^ (6 weeks): mean ± SEM = 7.4 ± 0.45 viable pups, n=35 litters from 11 females with 2-5 litters each; Na_v_1.8^LKB1^ (6 weeks): mean ± SEM = 12.1 ± 0.8 viable pups, n=31 litters from 11 females with 2-5 litters each; LKB1^fl/fl^ (20 weeks): mean ± SEM = 5.7 ± 1.022, n=6 litters from 3 females with 2 litters each; Na_v_1.8^LKB1^ (20 weeks): mean ± SEM = 7.4 ± 0.85 viable pups, n=10 litters from 5 females with 2 litters each. *p<0.05, **p<0.01, ***p<0.0001, ****p<0.0001 by ordinary one-way ANOVA (C-F) or ordinary two-way ANOVA (G).

**Supplementary Figure 2.**
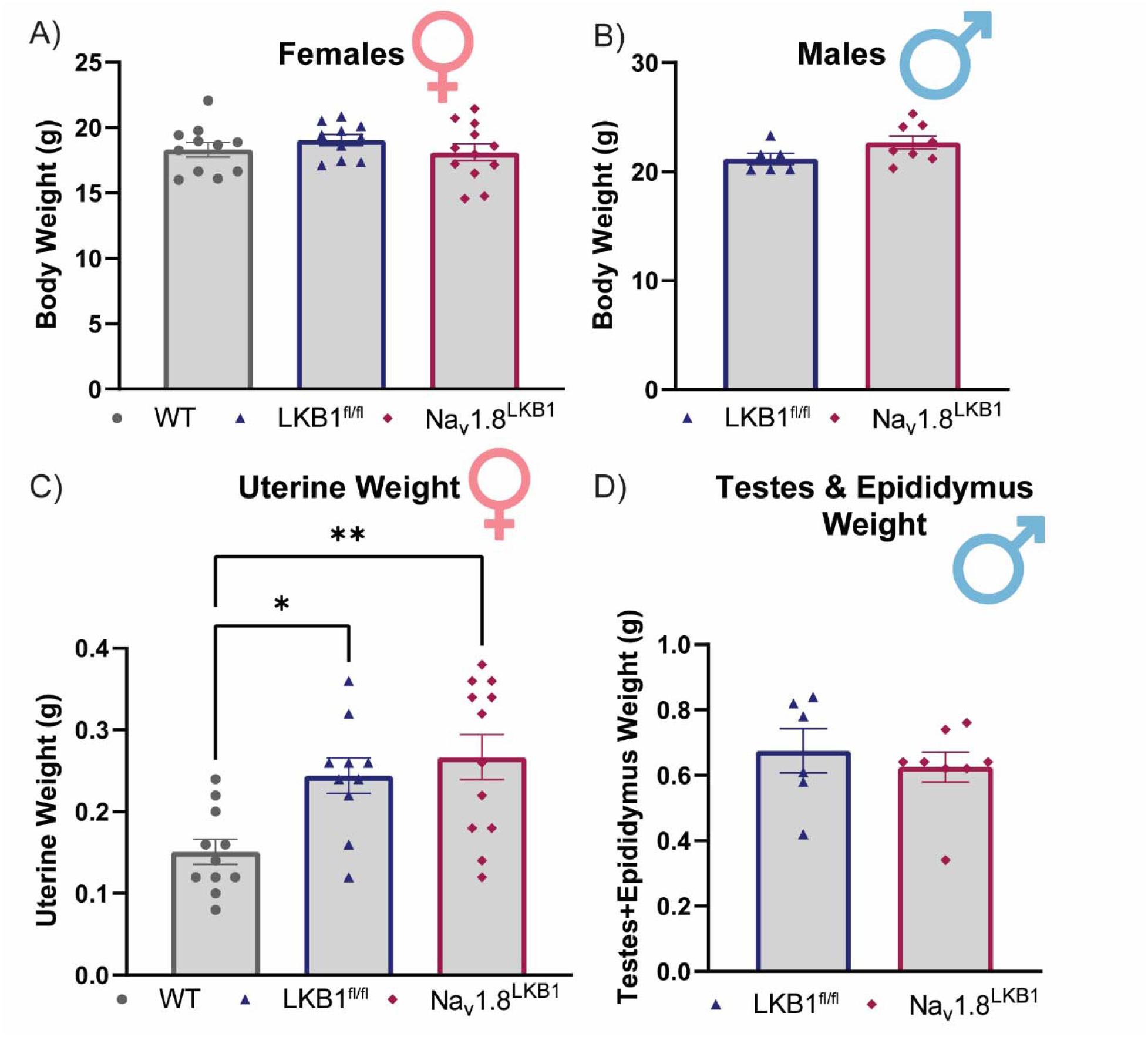
Histological analysis of reproductive organs in Nav1.8LKB1 males and females. (A) Body weight of female mice at six weeks of age. (WT: mean ± SEM = 18.3 ± 0.56 g, n=11 females; LKB1^fl/fl^: mean ± SEM = 19.1 ± 0.43 g, n=10 females; Na_v_1.8^LKB1^: mean ± SEM = 18.1 ± 0.63, n=12 females). (B) Body weight of male mice at six weeks of age. (LKB1^fl/fl^ mean ± SEM = 21.2 ± 0.51 g, n=6 males; Na_v_1.8^LKB1^: mean ± SEM = 22.7 ± 0.61 g, n=8 males). (C) Uterine weight of female mice at six weeks of age. (WT: mean ± SEM = 0.15 ± 0.015 g, n=11 females; LKB1^fl/fl^: mean ± SEM = 0.24 ± 0.022 g, n=10 females; Na_v_1.8^LKB1^: mean ± SEM = 0.27 ± 0.027, n=12 females). (D) Testes and epididymis weight of male mice at six weeks of age. (LKB1^fl/fl^: mean ± SEM = 0.68 ± 0.068 g, n=6 males; Na_v_1.8^LKB1^: mean ± SEM = 0.63 ± 0.045 g, n=8 males). *p<0.05, **p<0.01 by ordinary one-Way ANOVA (A, C) or unpaired two-tailed t-test (B, D).

**Supplementary Figure 3.**
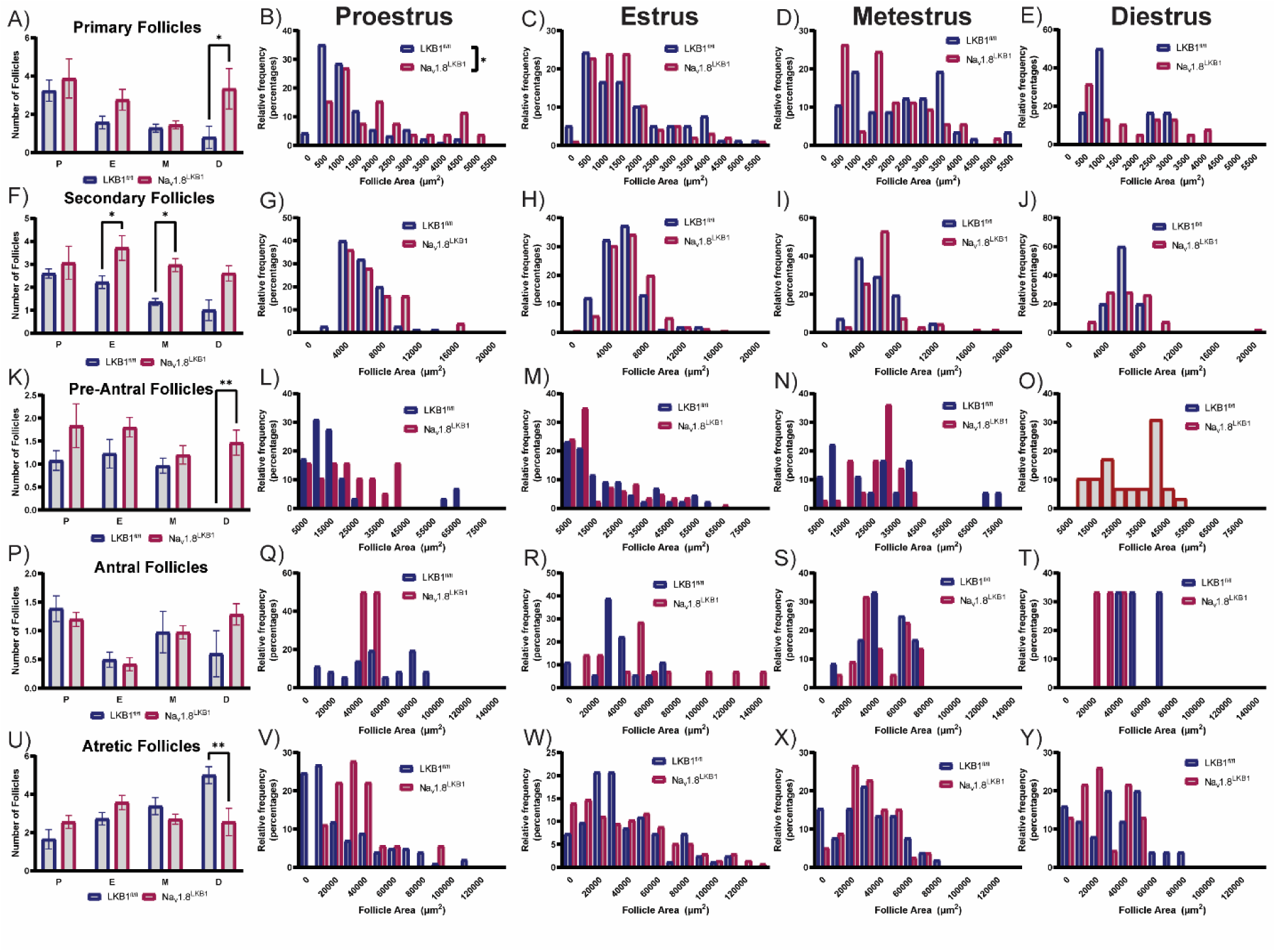
Follicular dynamics across the estrous cycle in Na_v_1.8^LKB1^ ovaries. (A) Number of primary follicles per ovary in each phase of the estrous cycle. Size of primary follicles in proestrus (B), estrus (C), metestrus (D), and diestrus (E). (F) Number of secondary follicles per ovary in each phase of the estrous cycle. Size of secondary follicles in proestrus (G), estrus (H), metestrus (I), and diestrus (J). (K) Number of pre-antral follicles per ovary in each phase of the estrous cycle. Size of pre-antral follicles in proestrus (L), estrus (M), metestrus (N), and diestrus (O). (P) Number of antral follicles per ovary in each phase of the estrous cycle. Size of antral follicles in proestrus (Q), estrus (R), metestrus (S), and diestrus (T). (U) Number of atretic follicles per ovary in each phase of the estrous cycle. Size of atretic follicles in proestrus (V), estrus (W), metestrus (X), and diestrus (Y). Size of all follicle area histograms is presented in µm^2^. P=proestrus, E=estrus, M=metestrus, D=diestrus. n=3-5 females/phase/genotype. Individual follicles were considered as technical replicates. \**p*<0.05, \*\**p*<0.01 by ordinary two-way ANOVA (phase x genotype). Data for (A, F, K, P, U) are represented as mean ± SEM. Data for (B-E, G-J, L-O, Q-T, and V-Y) are represented as histograms with each cycle phase on a separate graph.

**Supplementary Figure 4.**
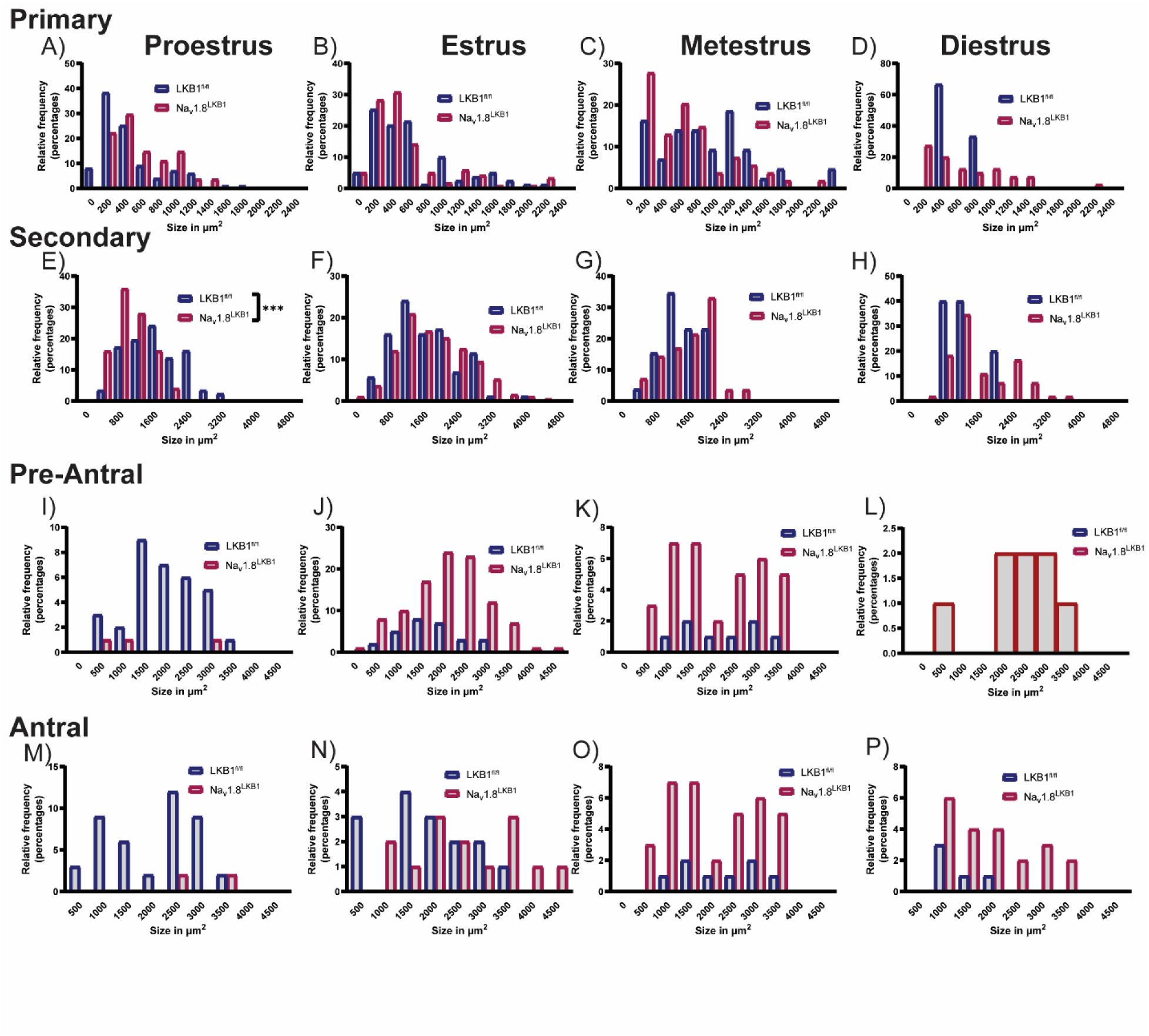
Oocyte growth across the estrous cycle. Size of oocytes from primary follicles in proestrus (A), estrus (B), metestrus (C), and diestrus (D). Size of oocytes from secondary follicles in proestrus (E), estrus (F), metestrus (G), and diestrus (H). Size of oocytes from pre-antral follicles in proestrus (I), estrus (J), metestrus (K), and diestrus (L). Size of oocytes from primary follicles in proestrus (M), estrus (N), metestrus (O), and diestrus (P). Size of all oocyte area histograms is presented in µm^2^. n=3-5 females/phase/genotype. Individual follicles were considered technical replicates. ***p<0.001 by Ordinary Two-Way ANOVA (phase x genotype). Data are represented as histograms with each cycle phase on a separate graph.

**Supplementary Table 1.** Statistics corresponding to Supplementary Figure 1. Litter Sizes, age at first litter, days to first litter, and days between litters were analyzed using ordinary one-eay ANOVA with *post hoc* Tukey’s multiple comparisons. Litter dize by fam age was analyzed using ordinary two-way ANOVA with *post hoc* Tukey’s multiple comparisons. Significance was set at *p<0.05* for all datasets. All significant results are in bold.

| Figure | Dataset | Main Effect |  | Multiple Comparisons |  |
| --- | --- | --- | --- | --- | --- |
|  |  | F (DFn, DFd) | p-value | Groups | p-value |
| S1C | Litter Sizes | F (4, 144) = 20.63 | **p<0.0001** | WT (C57) vs WT (mix) | p=0.6101 |
|  |  |  |  | WT (C57) vs LKB1<sup>fl/fl</sup> | p=0.4710 |
|  |  |  |  | **WT (C57) vs Nav1.8<sup>LKB1</sup>** | **p<0.0001** |
|  |  |  |  | **WT (C57) vs ICR (Outbred)** | **p<0.0001** |
|  |  |  |  | WT (mix) vs LKB1<sup>fl/fl</sup> | p=0.9996 |
|  |  |  |  | **WT (mix) vs Nav1.8LKB1** | **p<0.0001** |
|  |  |  |  | **WT (mix) vs ICR (Outbred)** | **p<0.0001** |
|  |  |  |  | LKB1<sup>fl/fl</sup> vs Nav1.8<sup>LKB1</sup> | **p<0.0001** |
|  |  |  |  | LKB1<sup>fl/fl</sup> vs ICR (Outbred) | **p<0.0001** |
|  |  |  |  | Nav1.8<sup>LKB1</sup> vs ICR (Outbred) | p=0.9398 |
| S1D | Age at First Litter | F (4, 38) = 5.475 | **p=0.0014** | WT (C57) vs WT (mix) | p=0.1545 |
|  |  |  |  | WT (C57) vs LKB1<sup>fl/fl</sup> | p=0.9996 |
|  |  |  |  | WT (C57) vs Nav1.8<sup>LKB1</sup> | p=0.0955 |
|  |  |  |  | WT (C57) vs ICR (Outbred) | p=0.9795 |
|  |  |  |  | WT (mix) vs LKB1<sup>fl/fl</sup> | p=0.0998 |
|  |  |  |  | **WT (mix) vs Nav1.8LKB1** | **p=0.0004** |
|  |  |  |  | WT (mix) vs ICR (Outbred) | p=0.1105 |
|  |  |  |  | LKB1<sup>fl/fl</sup> vs Nav1.8<sup>LKB1</sup> | p=0.1285 |
|  |  |  |  | LKB1<sup>fl/fl</sup> vs ICR (Outbred) | p=0.9938 |
|  |  |  |  | Nav1.8<sup>LKB1</sup> vs ICR (Outbred) | p=0.5605 |
| S1E | Days to First Litter | F (4, 32) = 1.926 | p=0.1301 | n/a |  |
| S1F | Days between Litters | F (4, 83) = 11.94 | **p<0.0001** | WT (C57) vs WT (mix) | p=0.8958 |
|  |  |  |  | WT (C57) vs LKB1<sup>fl/fl</sup> | p=0.2472 |
|  |  |  |  | WT (C57) vs Nav1.8<sup>LKB1</sup> | p=0.4498 |
|  |  |  |  | **WT (C57) vs ICR (Outbred)** | **p=0.0016** |
|  |  |  |  | WT (mix) vs LKB1<sup>fl/fl</sup> | p=0.8195 |
|  |  |  |  | WT (mix) vs Nav1.8<sup>LKB1</sup> | p=0.9650 |
|  |  |  |  | **WT (mix) vs ICR (Outbred)** | **p<0.0001** |
|  |  |  |  | LKB1<sup>fl/fl</sup> vs Nav1.8<sup>LKB1</sup> | p=0.9850 |
|  |  |  |  | LKB1<sup>fl/fl</sup> vs ICR (Outbred) | **p<0.0001** |
|  |  |  |  | **Nav1.8<sup>LKB1</sup> vs ICR (Outbred)** | **p<0.0001** |
| S1G | Litter Size by Dam Age | Genotype: F (1, 82) = 9.792 | **p=0.0024** | **6 weeks: LKB1<sup>fl/fl</sup> vs 6 weeks Nav1.8<sup>LKB1</sup>** | **p<0.0001** |
|  |  | Age: F (1, 82) = 10.06 | **p=0.0021** | 6 weeks: LKB1<sup>fl/fl</sup> vs 20 weeks: LKB1<sup>fl/fl</sup> | p=0.6785 |
|  |  | Genotype x Age: F (1, 82) = 2.065 | p=0.1545 | 6 weeks: LKB1<sup>fl/fl</sup> vs 20 weeks: Nav1.8<sup>LKB1</sup> | p>0.9999 |
|  |  |  |  | **6 weeks: Na<sub>v</sub>1.8<sup>LKB1</sup> vs 20 weeks: LKB1<sup>fl/fl</sup>** | ***p*=0.0007** |
|  |  |  |  | **6 weeks: Na<sub>v</sub>1.8<sup>LKB1</sup> vs 20 weeks: Na<sub>v</sub>1.8<sup>LKB1</sup>** | ***p*=0.0026** |
|  |  |  |  | 20 weeks: LKB1<sup>fl/fl</sup> vs 20 weeks: Na<sub>v</sub>1.8<sup>LKB1</sup> | *p*=0.7875 |

**Supplementary Table 2.**
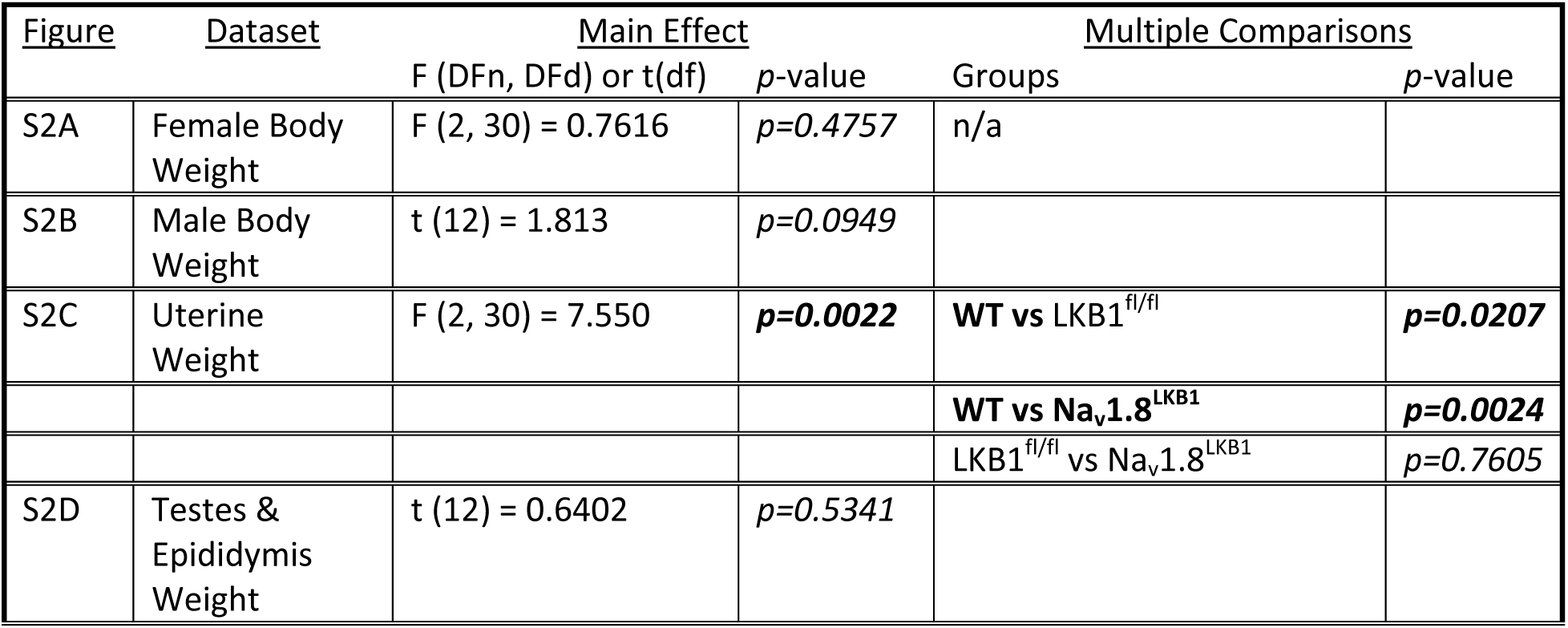
Statistics corresponding to Supplementary Figure 2. Data for female body weight and uterine weight were analyzed using ordinary one-way ANOVA with *post hoc* Tukey’s multiple comparisons. Data for male body weight and testes & epididymis weight were analyzed using unpaired two-tailed t-tests. Significance was set at *p<0.05* for all tests. Significant results are in bold.

| Figure | Dataset | Main Effect |  | Multiple Comparisons |  |
| --- | --- | --- | --- | --- | --- |
|  |  | F (DFn, DFd) or t(df) | p-value | Groups | p-value |
| S2A | Female Body Weight | F (2, 30) = 0.7616 | *p*=0.4757 | n/a |  |
| S2B | Male Body Weight | t (12) = 1.813 | *p*=0.0949 |  |  |
| S2C | Uterine Weight | F (2, 30) = 7.550 | ***p*=0.0022** | **WT vs LKB1<sup>fl/fl</sup>** | ***p*=0.0207** |
|  |  |  |  | **WT vs Na<sub>v</sub>1.8<sup>LKB1</sup>** | ***p*=0.0024** |
|  |  |  |  | **LKB1<sup>fl/fl</sup> vs Na<sub>v</sub>1.8<sup>LKB1</sup>** | ***p*=0.7605** |
| S2D | Testes & Epididymis Weight | t (12) = 0.6402 | *p*=0.5341 |  |  |

**Supplementary Table 3.** Statistics corresponding to Supplementary Figure 3. All datasets were analyzed using ordinary two-way ANOVA with *post hoc* Sidak’s multiple comparisons. Figures S3L-O (size of pre-antral follicles) was analyzed using ordinary two-way ANOVA with main effects only, as there were no pre-antral follicles observed in LKB1^fl/fl^ females during diestrus. Significance was set at *p<0.05* for all datasets. Significant results are in bold.

| Figure | Dataset | Main Effect |  | Multiple Comparisons |  |
| --- | --- | --- | --- | --- | --- |
|  |  | F (DFn, DFd) | p-value | Groups | p-value |
| S3A | Number of Primary Follicles | Phase: F (3, 21) = 4.783 | **p=0.0108** | Proestrus: LKB1<sup>fl/fl</sup> vs Na<sub>v</sub>1.8<sup>LKB1</sup> | p=0.9128 |
|  |  | Genotype: F (1, 21) = 7.419 | **p=0.0127** | Estrus: LKB1<sup>fl/fl</sup> vs Na<sub>v</sub>1.8<sup>LKB1</sup> | p=0.4073 |
|  |  | Phase x Genotype: F (3, 21) = 1.372 | p=0.2786 | Metestrus: LKB1<sup>fl/fl</sup> vs Na<sub>v</sub>1.8<sup>LKB1</sup> | p=0.9994 |
|  |  |  |  | **Diestrus: LKB1<sup>fl/fl</sup> vs Na<sub>v</sub>1.8<sup>LKB1</sup>** | **p=0.0399** |
| S3B-E | Size of Primary Follicles | Phase: F (3, 21) = 1.394 | p=0.2724 | **Proestrus: LKB1<sup>fl/fl</sup> vs Na<sub>v</sub>1.8<sup>LKB1</sup>** | **p=0.0374** |
|  |  | Genotype: F (1, 21) = 1.076 | p=0.3114 | Estrus: LKB1<sup>fl/fl</sup> vs Na<sub>v</sub>1.8<sup>LKB1</sup> | p=0.9899 |
|  |  | Phase x Genotype: F (3, 21) = 3.646 | **p=0.0293** | Metestrus: LKB1<sup>fl/fl</sup> vs Na<sub>v</sub>1.8<sup>LKB1</sup> | p=0.4533 |
|  |  |  |  | Diestrus: LKB1<sup>fl/fl</sup> vs Na<sub>v</sub>1.8<sup>LKB1</sup> | p=0.7835 |
| S3F | Number of Secondary Follicles | Phase: F (3, 21) = 3.973 | **p=0.0218** | Proestrus: LKB1<sup>fl/fl</sup> vs Na<sub>v</sub>1.8<sup>LKB1</sup> | p=0.8834 |
|  |  | Genotype: F (1, 21) = 22.02 | **p=0.0001** | Estrus: LKB1<sup>fl/fl</sup> vs Na<sub>v</sub>1.8<sup>LKB1</sup> | **p=0.0245** |
|  |  | Phase x Genotype: F (3, 21) = 0.9964 | p=0.4138 | **Metestrus: LKB1<sup>fl/fl</sup> vs Na<sub>v</sub>1.8<sup>LKB1</sup>** | **p=0.0345** |
|  |  |  |  | Diestrus: LKB1<sup>fl/fl</sup> vs Na<sub>v</sub>1.8<sup>LKB1</sup> | p=0.0523 |
| S3G-J | Size of Secondary Follicles | Phase: F (3, 21) = 1.386 | p=0.2746 | n/a |  |
|  |  | Genotype: F (1, 21) = 0.1568 | p=0.6962 |  |  |
|  |  | Phase x Genotype: F (3, 21) = 1.093 | p=0.3740 |  |  |
| S3K | Number of Pre-Antral Follicles | Phase: F (3, 21) = 3.490 | **p=0.0338** | Proestrus: LKB1<sup>fl/fl</sup> vs Na<sub>v</sub>1.8<sup>LKB1</sup> | p=0.2378 |
|  |  | Genotype: F (1, 21) = 15.62 | **p=0.0007** | Estrus: LKB1<sup>fl/fl</sup> vs Na<sub>v</sub>1.8<sup>LKB1</sup> | p=0.3598 |
|  |  | Phase x Genotype: F (3, 21) = 1.665 | p=0.2050 | Metestrus: LKB1<sup>fl/fl</sup> vs Na<sub>v</sub>1.8<sup>LKB1</sup> | p=0.9589 |
|  |  |  |  | **Diestrus: LKB1<sup>fl/fl</sup> vs Na<sub>v</sub>1.8<sup>LKB1</sup>** | **p=0.0077** |
| S3L-O | Size of Pre- | Phase: F (3, 21) | p=0.1100 | n/a |  |
|  | Antral Follicles | = 2.271 |  |  |  |
|  |  | Genotype: F (1, 21) = 0.1816 | *p*=0.6743 |  |  |
| S3P | Number of Antral Follicles | Phase: F (3, 21) = 4.982 | ***p*=0.0091** | Proestrus: LKB1<sup>fl/fl</sup> vs Na<sub>v</sub>1.8<sup>LKB1</sup> | *p*=0.9702 |
|  |  | Genotype: F (1, 21) = 4.049 | *p*=0.5314 | Estrus: LKB1<sup>fl/fl</sup> vs Na<sub>v</sub>1.8<sup>LKB1</sup> | *p*=0.9979 |
|  |  | Phase x Genotype: F (3, 21) = 1.295 | *p*=0.3022 | Metestrus: LKB1<sup>fl/fl</sup> vs Na<sub>v</sub>1.8<sup>LKB1</sup> | *p*>0.9999 |
|  |  |  |  | Diestrus: LKB1<sup>fl/fl</sup> vs Na<sub>v</sub>1.8<sup>LKB1</sup> | *p*=0.2447 |
| S3Q-T | Size of Antral Follicles | Phase: F (3, 21) = 0.6395 | *p*=0.5980 | Proestrus: LKB1<sup>fl/fl</sup> vs Na<sub>v</sub>1.8<sup>LKB1</sup> | *p*=0.9862 |
|  |  | Genotype: F (1, 21) = 0.08538 | *p*=0.7730 | Estrus: LKB1<sup>fl/fl</sup> vs Na<sub>v</sub>1.8<sup>LKB1</sup> | *p*=0.1104 |
|  |  | Phase x Genotype: F (3, 21) = 3.119 | ***p*=0.0478** | Metestrus: LKB1<sup>fl/fl</sup> vs Na<sub>v</sub>1.8<sup>LKB1</sup> | *p*=0.9998 |
|  |  |  |  | Diestrus: LKB1<sup>fl/fl</sup> vs Na<sub>v</sub>1.8<sup>LKB1</sup> | *p*=0.2517 |
| S3U | Number of Atretic Follicles | Phase: F (3, 21) = 4.496 | ***p*=0.0138** | Proestrus: LKB1<sup>fl/fl</sup> vs Na<sub>v</sub>1.8<sup>LKB1</sup> | *p*=0.5230 |
|  |  | Genotype: F (1, 21) = 1.172 | *p*=0.2913 | Estrus: LKB1<sup>fl/fl</sup> vs Na<sub>v</sub>1.8<sup>LKB1</sup> | *p*=0.4594 |
|  |  | Phase x Genotype: F (3, 21) = 6.006 | ***p*=0.0040** | Metestrus: LKB1<sup>fl/fl</sup> vs Na<sub>v</sub>1.8<sup>LKB1</sup> | *p*=0.7606 |
|  |  |  |  | **Diestrus: LKB1<sup>fl/fl</sup> vs Na<sub>v</sub>1.8<sup>LKB1</sup>** | ***p*=0.0068** |
| S3V-Y | Size of Atretic Follicles | Phase: F (3, 21) = 7.544 | ***p*=0.0013** | Proestrus: LKB1<sup>fl/fl</sup> vs Na<sub>v</sub>1.8<sup>LKB1</sup> | *p*=0.3232 |
|  |  | Genotype: F (1, 21) = 0.3060 | *p*=0.5860 | Estrus: LKB1<sup>fl/fl</sup> vs Na<sub>v</sub>1.8<sup>LKB1</sup> | *p*=0.9577 |
|  |  | Phase x Genotype: F (3, 21) = 1.787 | *p*=0.1806 | Metestrus: LKB1<sup>fl/fl</sup> vs Na<sub>v</sub>1.8<sup>LKB1</sup> | *p*=0.9915 |
|  |  |  |  | Diestrus: LKB1<sup>fl/fl</sup> vs Na<sub>v</sub>1.8<sup>LKB1</sup> | *p*=0.4770 |

**Supplementary Table 4.** Statistics corresponding to Supplementary Figure 4. All datasets were analyzed using ordinary two-way ANOVA with *post hoc* Sidak’s multiple comparisons. Figures S4I-L (size of pre-antral oocytes) was analyzed using ordinary two-way ANOVA with main effects only, as there were no pre-antral follicles observed in LKB1^fl/fl^ females during diestrus. Significance was set at *p<0.05* for all datasets. Significant results are in bold.

| Figure | Dataset | Main Effect |  | Multiple Comparisons |  |
| --- | --- | --- | --- | --- | --- |
|  |  | F (DFn, DFd) | p-value | Groups | p-value |
| S4A-D | Size of Primary Oocytes | Phase: F (3, 21) = 5.431 | **p=0.0063** | Proestrus: LKB1<sup>fl/fl</sup> vs Na<sub>v</sub>1.8<sup>LKB1</sup> | p=0.6163 |
|  |  | Genotype: F (1, 21) = 0.04228 | p=0.8391 | Estrus: LKB1<sup>fl/fl</sup> vs Na<sub>v</sub>1.8<sup>LKB1</sup> | p=0.9570 |
|  |  | Phase x Genotype: F (3, 21) = 3.478 | **p=0.0341** | Metestrus: LKB1<sup>fl/fl</sup> vs Na<sub>v</sub>1.8<sup>LKB1</sup> | p=0.0663 |
|  |  |  |  | Diestrus: LKB1<sup>fl/fl</sup> vs Na<sub>v</sub>1.8<sup>LKB1</sup> | p=0.5734 |
| S4E-H | Size of Secondary Oocytes | Phase: F (3, 21) = 9.431 | **p=0.0004** | **Proestrus: LKB1<sup>fl/fl</sup> vs Na<sub>v</sub>1.8<sup>LKB1</sup>** | **p=0.0033** |
|  |  | Genotype: F (1, 21) = 0.7770 | p=0.3880 | Estrus: LKB1<sup>fl/fl</sup> vs Na<sub>v</sub>1.8<sup>LKB1</sup> | p=0.2736 |
|  |  | Phase x Genotype: F (3, 21) = 9.009 | **p=0.0005** | Metestrus: LKB1<sup>fl/fl</sup> vs Na<sub>v</sub>1.8<sup>LKB1</sup> | p=0.8299 |
|  |  |  |  | **Diestrus: LKB1<sup>fl/fl</sup> vs Na<sub>v</sub>1.8<sup>LKB1</sup>** | **p=0.0358** |
| S4I-L | Size of Pre-Antral Oocytes | Phase: F (3, 21) = 1.682 | p=0.2013 | n/a |  |
|  |  | Genotype: F (1, 21) = 0.2454 | p=0.6255 |  |  |
| S4M-P | Size of Antral Oocytes | Phase: F (3, 21) = 4.446 | **p=0.0144** | Proestrus: LKB1<sup>fl/fl</sup> vs Na<sub>v</sub>1.8<sup>LKB1</sup> | p=0.1413 |
|  |  | Genotype: F (1, 21) = 7.916 | **p=0.0104** | Estrus: LKB1<sup>fl/fl</sup> vs Na<sub>v</sub>1.8<sup>LKB1</sup> | p=0.3491 |
|  |  | Phase x Genotype: F (3, 21) = 0.7569 | p=0.5307 | Metestrus: LKB1<sup>fl/fl</sup> vs Na<sub>v</sub>1.8<sup>LKB1</sup> | p=0.3346 |
|  |  |  |  | Diestrus: LKB1<sup>fl/fl</sup> vs Na<sub>v</sub>1.8<sup>LKB1</sup> | p>0.9999 |

## References

Agarwal A, Mulgund A, Hamada A, Chyatte MR (2015) A unique view on male infertility around the globe. Reproductive Biology and Endocrinology 13: 37

Alessi DR, Sakamoto K, Bayascas JR (2006) LKB1-Dependent Signaling Pathways. Annual Review of Biochemistry 75: 137–163

Arnal J-F, Lenfant F, Metivier R, Flouriot G, Henrion D, Adlanmerini M, Fontaine C, Gourdy P, Chambon P, Katzenellenbogen B et al (2017) Membrane and Nuclear Estrogen Receptor Alpha Actions: From Tissue Specificity to Medical Implications. Physiological Reviews 97: 1045–1087

Arnold J, Barcena de Arellano ML, Rüster C, Vercellino GF, Chiantera V, Schneider A, Mechsner S (2012) Imbalance between sympathetic and sensory innervation in peritoneal endometriosis. Brain, Behavior, and Immunity 26: 132–141

Asada N, Sanada K, Fukada Y (2007) LKB1 Regulates Neuronal Migration and Neuronal Differentiation in the Developing Neocortex through Centrosomal Positioning. The Journal of Neuroscience 27: 11769

Bardeesy N, Sinha M, Hezel AF, Signoretti S, Hathaway NA, Sharpless NE, Loda M, Carrasco DR, DePinho RA (2002) Loss of the Lkb1 tumour suppressor provokes intestinal polyposis but resistance to transformation. Nature 419: 162–167

Barnes AP, Lilley BN, Pan YA, Plummer LJ, Powell AW, Raines AN, Sanes JR, Polleux F (2007) LKB1 and SAD Kinases Define a Pathway Required for the Polarization of Cortical Neurons. Cell 129: 549–563

Brabaharan S, Veettil SK, Kaiser JE, Raja Rao VR, Wattanayingcharoenchai R, Maharajan M, Insin P, Talungchit P, Anothaisintawee T, Thakkinstian A et al (2022) Association of Hormonal Contraceptive Use With Adverse Health Outcomes: An Umbrella Review of Meta-analyses of Randomized Clinical Trials and Cohort Studies. JAMA Network Open 5: e2143730–e2143730

Brown K, Brown K, Simpson E, Simpson E (2009) Metformin Inhibits Aromatase Expression in Primary Human Breast Adipose Stromal Cells. Cancer Research 69: 3132–3132

Burden HW, Leonard M, Smith CP, Lawrence Jr IE (1983) The sensory innervation of the ovary: A horseradish peroxidase study in the rat. The Anatomical Record 207: 623–627

Burger CA, Alevy J, Casasent AK, Jiang D, Albrecht NE, Liang JH, Hirano AA, Brecha NC, Samuel MA (2020) LKB1 coordinates neurite remodeling to drive synapse layer emergence in the outer retina. eLife 9: e56931

Byers SL, Wiles MV, Dunn SL, Taft RA (2012) Mouse estrous cycle identification tool and images. PLoS One 7: e35538

Calka J, McDonald JK, Ojeda SR (1988) The Innervation of the Immature Rat Ovary by Calcitonin Gene-Related Peptide1. Biology of Reproduction 39: 1215–1223

Carson SA, Kallen AN (2021) Diagnosis and Management of Infertility: A Review. JAMA 326: 65–76

Chaban V, Christensen A, Wakamatsu M, McDonald M, Rapkin A, McDonald J, Micevych P (2007) The same dorsal root ganglion neurons innervate uterus and colon in the rat. NeuroReport 18

Chen IC, Chang Y-C, Lu Y-S, Chung K-P, Huang C-S, Lu T-P, Kuo W-H, Wang M-Y, Kuo K-T, Wu P-F et al (2016) Clinical Relevance of Liver Kinase B1(LKB1) Protein and Gene Expression in Breast Cancer. Scientific Reports 6: 21374

Chen O, Donnelly CR, Ji R-R (2020) Regulation of pain by neuro-immune interactions between macrophages and nociceptor sensory neurons. Current Opinion in Neurobiology 62: 17–25

Chiu IM, Heesters BA, Ghasemlou N, Von Hehn CA, Zhao F, Tran J, Wainger B, Strominger A, Muralidharan S, Horswill AR et al (2013) Bacteria activate sensory neurons that modulate pain and inflammation. Nature 501: 52–57

Christensen A, Bentley GE, Cabrera R, Ortega HH, Perfito N, Wu TJ, Micevych P (2012) Hormonal Regulation of Female Reproduction. Horm Metab Res 44: 587–591

Colledge WH (2013) The neuroendocrine regulation of the mammalian reproductive axis. Introduction. Exp Physiol 98: 1519–1521

Cora MC, Kooistra L, Travlos G (2015) Vaginal Cytology of the Laboratory Rat and Mouse: Review and Criteria for the Staging of the Estrous Cycle Using Stained Vaginal Smears. Toxicol Pathol 43: 776–793

Courchet J, Lewis Tommy L, Jr., Lee S, Courchet V, Liou D-Y, Aizawa S, Polleux F (2013) Terminal Axon Branching Is Regulated by the LKB1-NUAK1 Kinase Pathway via Presynaptic Mitochondrial Capture. Cell 153: 1510–1525

Cruz G, Fernandois D, Paredes AH (2017) Ovarian function and reproductive senescence in the rat: role of ovarian sympathetic innervation. Reproduction 153: R59–R68

Dahlman-Wright K, Cavailles V, Fuqua SA, Jordan VC, Katzenellenbogen JA, Korach KS, Maggi A, Muramatsu M, Parker MG, Gustafsson J-Å (2006) International Union of Pharmacology. LXIV. Estrogen Receptors. Pharmacological Reviews 58: 773

del Campo M, Piquer B, Witherington J, Sridhar A, Lara HE (2019) Effect of Superior Ovarian Nerve and Plexus Nerve Sympathetic Denervation on Ovarian-Derived Infertility Provoked by Estradiol Exposure to Rats. Frontiers in Physiology 10

Deng Y, Zhang Y, Li S, Zhou W, Ye L, Wang L, Tao T, Gu J, Yang Z, Zhao D et al (2017) Steroid hormone profiling in obese and nonobese women with polycystic ovary syndrome. Scientific Reports 7: 14156

Dodds KN, Kyloh MA, Travis L, Beckett EAH, Spencer NJ (2021) Morphological identification of thoracolumbar spinal afferent nerve endings in mouse uterus. Journal of Comparative Neurology 529: 2029–2041

dos Santos NL, Lenert ME, Castillo ZW, Mody PH, Thompson LT, Burton MD (2022) Age and Sex Drive Differential Behavioral and Neuroimmune Phenotypes During Postoperative Pain. Neurobiology of Aging

Engler-Chiurazzi EB, Chastain WH, Citron KK, Lambert LE, Kikkeri DN, Shrestha SS (2022) Estrogen, the Peripheral Immune System and Major Depression – A Reproductive Lifespan Perspective. Frontiers in Behavioral Neuroscience 16

Fauser BCJM, van Heusden AM (1997) Manipulation of Human Ovarian Function: Physiological Concepts and Clinical Consequences*. Endocrine Reviews 18: 71–106

Flores A, Velasco J, Gallegos AI, Mendoza FD, Everardo PM, Cruz M-E, Domínguez R (2011) Acute effects of unilateral sectioning the superior ovarian nerve of rats with unilateral ovariectomy on ovarian hormones (progesterone, testosterone and estradiol) levels vary during the estrous cycle. Reproductive Biology and Endocrinology 9: 34

Furat Rencber S, Kurnaz Ozbek S, Eraldemır C, Sezer Z, Kum T, Ceylan S, Guzel E (2018) Effect of resveratrol and metformin on ovarian reserve and ultrastructure in PCOS: an experimental study. Journal of Ovarian Research 11: 55

Garner KM, Burton MD (2022) Sex-specific role of sensory neuron LKB1 on metabolic stress-induced mechanical hypersensitivity and mitochondrial respiration. American Journal of Physiology-Regulatory, Integrative and Comparative Physiology 323: R227–R243

Gautron L, Sakata I, Udit S, Zigman JM, Wood JN, Elmquist JK (2011) Genetic tracing of Nav1.8-expressing vagal afferents in the mouse. Journal of Comparative Neurology 519: 3085–3101

Gaytan F, Morales C, Leon S, Heras V, Barroso A, Avendaño MS, Vazquez MJ, Castellano JM, Roa J, Tena-Sempere M (2017) Development and validation of a method for precise dating of female puberty in laboratory rodents: The puberty ovarian maturation score (Pub-Score). Scientific Reports 7: 46381

Gershon E, Dekel N (2020) Newly Identified Regulators of Ovarian Folliculogenesis and Ovulation. International Journal of Molecular Sciences 21: 4565

Guastella E, Longo RA, Carmina E (2010) Clinical and endocrine characteristics of the main polycystic ovary syndrome phenotypes. Fertil Steril 94: 2197–2201

Ham S, Brown KA, Simpson ER, Meachem SJ (2017) Immunolocalisation of aromatase regulators liver kinase B1, phosphorylated AMP-activated protein kinase and cAMP response element-binding protein-regulated transcription co-activators in the human testis. Reproduction, Fertility and Development 29: 1029–1038

Hanada T, Uchida S, Hotta H, Aikawa Y (2011) Number, size, conduction, and vasoconstrictor ability of unmyelinated fibers of the ovarian nerve in adult and aged rats. Autonomic Neuroscience: Basic and Clinical 164: 6–12

Herweijer G, Kyloh M, Beckett EAH, Dodds KN, Spencer NJ (2014) Characterization of primary afferent spinal innervation of mouse uterus. Frontiers in Neuroscience 8

Herzig S, Shaw RJ (2018) AMPK: guardian of metabolism and mitochondrial homeostasis. Nature Reviews Molecular Cell Biology 19: 121–135

Hohos NM, Cho KJ, Swindle DC, Skaznik-Wikiel ME (2018) High-fat diet exposure, regardless of induction of obesity, is associated with altered expression of genes critical to normal ovulatory function. Molecular and Cellular Endocrinology 470: 199–207

Hsueh AJW, Kawamura K, Cheng Y, Fauser BCJM (2015) Intraovarian Control of Early Folliculogenesis. Endocrine Reviews 36: 1–24

Huang S, Ziegler CGK, Austin J, Mannoun N, Vukovic M, Ordovas-Montanes J, Shalek AK, von Andrian UH (2021) Lymph nodes are innervated by a unique population of sensory neurons with immunomodulatory potential. Cell 184: 441–459.e425

Huang W, She L, Chang X-y, Yang R-r, Wang L, Ji H-b, Jiao J-w, Poo M-m (2014) Protein kinase LKB1 regulates polarized dendrite formation of adult hippocampal newborn neurons. Proceedings of the National Academy of Sciences 111: 469–474

Inyama CO, Wharton J, Su HC, Polak JM (1986) CGRP-immunoreactive nerves in the genitalia of the female rat originate from dorsal root ganglia T11-L3 and L6-S1: A combined immunocytochemical and retrograde tracing study. Neuroscience Letters 69: 13–18

Jia L, Lee S, Tierney JA, Elmquist JK, Burton MD, Gautron L (2021) TLR4 signaling selectively and directly promotes CGRP release from vagal afferents in the mouse. Eneuro 8

Kagitani F, Uchida S, Hotta H (2008) Effects of Electrical Stimulation of the Superior Ovarian Nerve and the Ovarian Plexus Nerve on the Ovarian Estradiol Secretion Rate in Rats. The Journal of Physiological Sciences 58: 133–138

Klein CM, Burden HW (1988) Substance P- and vasoactive intestinal polypeptide (VIP)-immunoreactive nerve fibers in relation to ovarian postganglionic perikarya in para- and prevertebral ganglia: Evidence from combined retrograde tracing and immunocytochemistry. Cell and Tissue Research 252: 403–410

Kocer D, Bayram F, Diri H (2014) The effects of metformin on endothelial dysfunction, lipid metabolism and oxidative stress in women with polycystic ovary syndrome. Gynecological Endocrinology 30: 367–371

Kupari J, Häring M, Agirre E, Castelo-Branco G, Ernfors P (2019) An Atlas of Vagal Sensory Neurons and Their Molecular Specialization. Cell Reports 27: 2508–2523.e2504

Kwon S-K, Sando R, III, Lewis TL, Hirabayashi Y, Maximov A, Polleux F (2016) LKB1 Regulates Mitochondria-Dependent Presynaptic Calcium Clearance and Neurotransmitter Release Properties at Excitatory Synapses along Cortical Axons. PLOS Biology 14: e1002516

Lainez NM, Coss D (2019) Obesity, Neuroinflammation, and Reproductive Function. Endocrinology 160: 2719–2736

Lainez NM, Jonak CR, Nair MG, Ethell IM, Wilson EH, Carson MJ, Coss D (2018) Diet-Induced Obesity Elicits Macrophage Infiltration and Reduction in Spine Density in the Hypothalami of Male but Not Female Mice. Frontiers in Immunology 9

Lara HE, Dorfman M, Venegas M, Luza SM, Luna SL, Mayerhofer A, Guimaraes MA, Rosa E Silva AAM, Ramírez VD (2002) Changes in sympathetic nerve activity of the mammalian ovary during a normal estrous cycle and in polycystic ovary syndrome: Studies on norepinephrine release. Microscopy Research and Technique 59: 495–502

Lawrence Jr IE, Burden HW (1980) The origin of the extrinsic adrenergic innervation to the rat ovary. The Anatomical Record 196: 51–59

Leitman DC, Paruthiyil S, Vivar OI, Saunier EF, Herber CB, Cohen I, Tagliaferri M, Speed TP (2010) Regulation of specific target genes and biological responses by estrogen receptor subtype agonists. Current Opinion in Pharmacology 10: 629–636

Lenert ME, Avona A, Garner KM, Barron LR, Burton MD (2021a) Sensory Neurons, Neuroimmunity, and Pain Modulation by Sex Hormones. Endocrinology 162

Lenert ME, Burton MD (2021) Acute effects of a high-fat diet on estrous cycling and body weight of intact female mice. Neuropsychopharmacology

Lenert ME, Chaparro MM, Burton MD (2021b) Homeostatic Regulation of Estrus Cycle of Young Female Mice on Western Diet. Journal of the Endocrine Society 5

Lim HJ, Wang H (2010) Uterine disorders and pregnancy complications: insights from mouse models. The Journal of Clinical Investigation 120: 1004–1015

Linher-Melville K, Zantinge S, Singh G (2012) Liver kinase B1 expression (LKB1) is repressed by estrogen receptor alpha (ERα) in MCF-7 human breast cancer cells. Biochemical and Biophysical Research Communications 417: 1063–1068

Lipovka Y, Chen H, Vagner J, Price Theodore J, Tsao T-S, Konhilas John P (2015) Oestrogen receptors interact with the α-catalytic subunit of AMP-activated protein kinase. Bioscience Reports 35

Lotti F, Marchiani S, Corona G, Maggi M (2021) Metabolic Syndrome and Reproduction. International Journal of Molecular Sciences 22

Magoffin DA (2005) Ovarian theca cell. The International Journal of Biochemistry & Cell Biology 37: 1344–1349

Magyar BP, Santi M, Sommer G, Nuoffer J-M, Leichtle A, Grössl M, Fluck CE (2022) Short-Term Fasting Attenuates Overall Steroid Hormone Biosynthesis in Healthy Young Women. Journal of the Endocrine Society 6

McInnes KJ, Brown KA, Hunger NI, Simpson ER (2012) Regulation of LKB1 expression by sex hormones in adipocytes. International Journal of Obesity 36: 982–985

Mikhael S, Punjala-Patel A, Gavrilova-Jordan L (2019) Hypothalamic-Pituitary-Ovarian Axis Disorders Impacting Female Fertility. Biomedicines 7

Miller MQ, Hernández IC, Chacko JV, Minderler S, Jowett N (2021) Two-photon excitation fluorescent spectral and decay properties of retrograde neuronal tracer Fluoro-Gold. Scientific Reports 11: 18053

Molina E, Hong L, Chefetz I (2021) AMPKα-like proteins as LKB1 downstream targets in cell physiology and cancer. Journal of Molecular Medicine 99: 651–662

Mondello SE, Jefferson SC, O’Steen WA, Howland DR (2016) Enhancing Fluorogold-based neural tract tracing. Journal of Neuroscience Methods 270: 85–91

Mónica Brauer M, Smith PG (2015) Estrogen and female reproductive tract innervation: Cellular and molecular mechanisms of autonomic neuroplasticity. Autonomic Neuroscience: Basic and Clinical 187: 1–17

Morales-Ledesma L, Linares R, Rosas G, Morán C, Chavira R, Cárdenas M, Domínguez R (2010) Unilateral sectioning of the superior ovarian nerve of rats with polycystic ovarian syndrome restores ovulation in the innervated ovary. Reproductive Biology and Endocrinology 8: 99

Morales-Ledesma L, Trujillo A, Apolonio J (2015) In the pubertal rat, the regulation of ovarian function involves the synergic participation of the sensory and sympathetic innervations that arrive at the gonad. Reproductive Biology and Endocrinology 13: 61

Morales L, Ricardo B, Bolaños A, Chavira R, Domínguez R (2007) Ipsilateral vagotomy to unilaterally ovariectomized pre-pubertal rats modifies compensatory ovarian responses. Reproductive Biology and Endocrinology 5: 24

Mørch LS, Skovlund CW, Hannaford PC, Iversen L, Fielding S, Lidegaard Ø (2017) Contemporary Hormonal Contraception and the Risk of Breast Cancer. New England Journal of Medicine 377: 2228–2239

Mounier R, Théret M, Arnold L, Cuvellier S, Bultot L, Göransson O, Sanz N, Ferry A, Sakamoto K, Foretz M et al (2013) AMPKα1 Regulates Macrophage Skewing at the Time of Resolution of Inflammation during Skeletal Muscle Regeneration. Cell Metabolism 18: 251–264

Nakano A, Takashima S (2012) LKB1 and AMP-activated protein kinase: regulators of cell polarity. Genes to Cells 17: 737–747

Nath-Sain S, Marignani PA (2009) LKB1 Catalytic Activity Contributes to Estrogen Receptor α Signaling. Molecular Biology of the Cell 20: 2785–2795

National Research Council Committee for the Update of the Guide for the C, Use of Laboratory A (2011) The National Academies Collection: Reports funded by National Institutes of Health. In: Guide for the Care and Use of Laboratory Animals, National Academies Press (US) Copyright © 2011, National Academy of Sciences.: Washington (DC)

Negrón AL, Radovick S (2020) High-Fat Diet Alters LH Secretion and Pulse Frequency in Female Mice in an Estrous Cycle-Dependent Manner. Endocrinology 161

Papka RE, Storey-Workley M, Shughrue PJ, Merchenthaler I, Collins JJ, Usip S, Saunders PTK, Shupnik M (2001) Estrogen receptor-α and -β immunoreactivity and mRNA in neurons of sensory and autonomic ganglia and spinal cord. Cell and Tissue Research 304: 193–214

Pastelín CF, Rosas NH, Morales-Ledesma L, Linares R, Domínguez R, Morán C (2017) Anatomical organization and neural pathways of the ovarian plexus nerve in rats. Journal of Ovarian Research 10: 18

Paterni I, Granchi C, Katzenellenbogen JA, Minutolo F (2014) Estrogen receptors alpha (ERα) and beta (ERβ): Subtype-selective ligands and clinical potential. Steroids 90: 13–29

Patil MJ, Hovhannisyan AH, Akopian AN (2018) Characteristics of sensory neuronal groups in CGRP-cre-ER reporter mice: Comparison to Nav1.8-cre, TRPV1-cre and TRPV1-GFP mouse lines. PLOS ONE 13: e0198601

Pedersen T, Peters H (1968) PROPOSAL FOR A CLASSIFICATION OF OOCYTES AND FOLLICLES IN THE MOUSE OVARY. Reproduction 17: 555–557

Percie du Sert N, Ahluwalia A, Alam S, Avey MT, Baker M, Browne WJ, Clark A, Cuthill IC, Dirnagl U, Emerson M et al (2020) Reporting animal research: Explanation and elaboration for the ARRIVE guidelines 2.0. PLOS Biology 18: e3000411

Perrigo G, Bronson FH (1983) Foraging effort, food intake, fat deposition and puberty in female mice. Biol Reprod 29: 455–463

Pinho-Ribeiro FA, Verri WA, Jr., Chiu IM (2017) Nociceptor Sensory Neuron-Immune Interactions in Pain and Inflammation. Trends Immunol 38: 5–19

Ramírez Hernández DA, Vieyra Valdez E, Rosas Gavilán G, Linares Culebro R, Espinoza Moreno JA, Chaparro Ortega A, Domínguez Casalá R, Morales-Ledesma L (2020) Role of the superior ovarian nerve in the regulation of follicular development and steroidogenesis in the morning of diestrus 1. Journal of Assisted Reproduction and Genetics 37: 1477–1488

Robakis T, Williams KE, Nutkiewicz L, Rasgon NL (2019) Hormonal Contraceptives and Mood: Review of the Literature and Implications for Future Research. Current Psychiatry Reports 21: 57

Robker RL, Hennebold JD, Russell DL (2018) Coordination of Ovulation and Oocyte Maturation: A Good Egg at the Right Time. Endocrinology 159: 3209–3218

Rosas G, Linares R, Ramírez DA, Vieyra E, Trujillo A, Domínguez R, Morales-Ledesma L (2018) The Neural Signals of the Superior Ovarian Nerve Modulate in an Asymmetric Way the Ovarian Steroidogenic Response to the Vasoactive Intestinal Peptide. Frontiers in Physiology 9

Ruddenklau A, Campbell RE (2019) Neuroendocrine Impairments of Polycystic Ovary Syndrome. Endocrinology 160: 2230–2242

Russell FA, King R, Smillie SJ, Kodji X, Brain SD (2014) Calcitonin Gene-Related Peptide: Physiology and Pathophysiology. Physiological Reviews 94: 1099–1142

Saikia R, Joseph J (2021) AMPK: a key regulator of energy stress and calcium-induced autophagy. Journal of Molecular Medicine 99: 1539–1551

Sapio MR, Vazquez FA, Loydpierson AJ, Maric D, Kim JJ, LaPaglia DM, Puhl HL, Lu VB, Ikeda SR, Mannes AJ et al (2020) Comparative Analysis of Dorsal Root, Nodose and Sympathetic Ganglia for the Development of New Analgesics. Frontiers in Neuroscience 14

Schmued LC, Fallon JH (1986) Fluoro-gold: a new fluorescent retrograde axonal tracer with numerous unique properties. Brain Research 377: 147–154

Shackelford DB, Shaw RJ (2009) The LKB1–AMPK pathway: metabolism and growth control in tumour suppression. Nature Reviews Cancer 9: 563–575

Shelly M, Cancedda L, Heilshorn S, Sumbre G, Poo M-m (2007) LKB1/STRAD Promotes Axon Initiation During Neuronal Polarization. Cell 129: 565–577

Sirotkin AV, Fabian D, Babelova Kubandova J, Vlckova R, Alwasel S, Harrath AH (2018) Body fat affects mouse reproduction, ovarian hormone release, and response to follicular stimulating hormone. Reprod Biol 18: 5–11

Stirling LC, Forlani G, Baker MD, Wood JN, Matthews EA, Dickenson AH, Nassar MA (2005) Nociceptor-specific gene deletion using heterozygous NaV1. 8-Cre recombinase mice. Pain 113: 27-36

Szabo-Pardi TA, Barron LR, Lenert ME, Burton MD (2021) Sensory Neuron TLR4 mediates the development of nerve-injury induced mechanical hypersensitivity in female mice. Brain, Behavior, and Immunity 97: 42–60

Tingåker BK, Irestedt L (2010) Changes in uterine innervation in pregnancy and during labour. Current Opinion in Anesthesiology 23

Tiosano D, Mears JA, Buchner DA (2019) Mitochondrial Dysfunction in Primary Ovarian Insufficiency. Endocrinology 160: 2353–2366

Trujillo A, Morales L, Domínguez R (2015) The effects of sensorial denervation on the ovarian function, by the local administration of capsaicin, depend on the day of the oestrous cycle when the treatment was performed. Endocrine 48: 321–328

Wang Y, Leung VH, Zhang Y, Nudell VS, Loud M, Servin-Vences MR, Yang D, Wang K, Moya-Garzon MD, Li VL et al (2022) The role of somatosensory innervation of adipose tissues. Nature 609: 569–574

Wassif WS, McLoughlin DM, Vincent RP, Conroy S, Russell GF, Taylor NF (2011) Steroid metabolism and excretion in severe anorexia nervosa: effects of refeeding. The American Journal of Clinical Nutrition 93: 911–917

Westwood FR (2008) The female rat reproductive cycle: a practical histological guide to staging. Toxicol Pathol 36: 375–384

Winckler B (2007) BDNF Instructs the Kinase LKB1 To Grow an Axon. Cell 129: 459–460

Witchel SF, Oberfield SE, Peña AS (2019) Polycystic Ovary Syndrome: Pathophysiology, Presentation, and Treatment With Emphasis on Adolescent Girls. Journal of the Endocrine Society 3: 1545–1573

Yang Z-m, Yang M-f, Yu W, Tao H-m (2019) Molecular mechanisms of estrogen receptor β-induced apoptosis and autophagy in tumors: implication for treating osteosarcoma. Journal of International Medical Research 47: 4644–4655

Zhang X, Lu B, Huang X, Xu H, Zhou C, Lin J (2010) Innervation of endometrium and myometrium in women with painful adenomyosis and uterine fibroids. Fertility and Sterility 94: 730–737

Zoubina EV, Smith PG (2001) Sympathetic hyperinnervation of the uterus in the estrogen receptor α knock-out mouse. Neuroscience 103: 237–244

